# Structural inventory of native ribosomal ABCE1-43S pre-initiation complexes

**DOI:** 10.1101/2020.07.09.194902

**Authors:** Hanna Kratzat, Timur Mackens-Kiani, Michael Ameismeier, Jingdong Cheng, Estelle Dacheux, Abdelkader Namane, Otto Berninghausen, Micheline Fromont-Racine, Thomas Becker, Roland Beckmann

## Abstract

In eukaryotic translation, the termination and recycling phases are linked to subsequent initiation by persistence of several factors. These comprise the large eIF3 complex, eIF3j (Hcr1 in yeast) and the ATP-binding cassette protein ABCE1 (Rli1 in yeast). The ATPase is mainly active as a recycling factor, but it can remain bound to the dissociated 40S subunit until formation of 43S pre-initiation complexes. However, its functional role and native architectural context remains largely enigmatic. Here, we present an architectural inventory of native yeast and human ABCE1-containing pre-initiation complexes by cryo-EM. We found that ABCE1 was mostly associated with early 43S but also later 48S phases of initiation. It directly interacted with eIF3j *via* its unique iron-sulfur cluster domain and adopted a novel hybrid conformation, which was ATPase-inhibited and stabilized by an unknown factor bound between the nucleotide binding sites. Moreover, the native human samples provided a near-complete molecular picture of the architecture and sophisticated interaction network of the 43S-bound eIF3 complex and also the eIF2 ternary complex containing the initiator tRNA.

## Introduction

Translation of an mRNA into a polypeptide sequence is a central cellular process, which is highly regulated and linked to other cellular processes like ribosome biogenesis, mRNA turnover, and ribosome quality control. Most decisive for translational efficiency and regulation is the initiation phase; however, in eukaryotes the individual phases of translation were found to be coupled, especially termination with ribosome recycling and a new round of initiation. Two prominent examples are the conserved multisubunit complex eIF3, which has been described as a factor functioning across the translation cycle (Valasek *et al*, 2017), as well as the ATP-binding cassette (ABC) ATPase ABCE1 (Rli1 in *Saccharomyces cerevisiae*), which was shown to enhance termination activity of the eRF1 release factor and which represents the key enzyme for ATP-dependent ribosome recycling (Pisarev *et al*, 2010; Shoemaker & Green, 2011). Moreover, ABCE1 was found associated with initiation factors (Chen *et al*, 2006; Dong *et al*, 2004) and as a part of eIF3-containing 43S or 48S (pre-)initiation complexes (Andersen & Leevers, 2007; Preis *et al*, 2014; Mancera-Martinez *et al*, 2017).

The ABCE1 ATPase consists of two nucleotide binding domains (NBDs) that are forming two nucleotide binding sites (NBSs) at their interface, as well as an essential iron-sulfur cluster domain (FeSD) at its N-terminus (Barthelme *et al*, 2007; Hopfner, 2016). ABCE1 binds the 80S ribosome during canonical stop codon-dependent termination or during rescue of stalled ribosomes and splits the 80S ribosomes into 40S and 60S small (SSU) and large (LSU) subunits, respectively. This recycling reaction requires an A site factor in the ribosome, either release factor eRF1 (after termination) or its homologue Pelota (Dom34 in *S.c*.; for ribosome rescue), in order to form part of the interaction network for ABCE1 (Becker *et al*, 2012; Brown *et al*, 2015; Preis *et al.*, 2014). ABCE1 binds these pre-splitting complexes in a semi-open state with respect to its NBSs. Splitting requires binding of ATP and site-occlusion to both NBS (Barthelme *et al*, 2011; Gouridis *et al*, 2019; Nurenberg-Goloub *et al*, 2018) and can be recapitulated *in vitro* (Becker *et al.*, 2012; Nurenberg-Goloub & Tampe, 2019; Pisareva *et al*, 2011; Shao *et al*, 2015; Shoemaker & Green, 2011). After *in vitro* splitting, ABCE1 was observed to remain bound to the 40S small subunit to form a post-splitting complex (PSC), in which the two NBDs are present in a closed, nucleotide-occluding state (Heuer *et al*, 2017; Kiosze-Becker *et al*, 2016; Nürenberg-Goloub *et al*, 2020). Therefore, it was assumed that *in vivo* as well, ABCE1 may remain bound to the 40S for a defined time span (Gerovac & Tampe, 2019) to prevent re-association of the LSU (Heuer *et al.*, 2017) or to coordinate assembly of initiation factors on the 40S subunit. However, a direct physical involvement of ABCE1 in the translation initiation process has not been shown to date.

In eukaryotes, the start of translation initiation requires the assembly of the 43S pre-initiation complex (PIC). It consists of the 40S subunit, eIF3, eIF1, eIF1A, eIF5, and the ternary complex (TC) formed by the trimeric eIF2-*αβγ*, initiator methionyl tRNA (tRNA_i_), and GTP. After 43S PIC assembly, the mRNA - in collaboration with the eIF4F complex (the cap-binding protein eIF4E, the helicase eIF4A and the scaffolding protein eIF4G) - can be recruited to the 43S PIC, forming the 48S initiation complex (IC). This event is coordinated by interactions between eIF3 and eIF4F as well as eIF4B, a single-stranded RNA-binding protein that attaches to the 40S subunit (Walker *et al*, 2013) and stimulates the helicase activity of eIF4A. The 48S complex then scans the mRNA for the first cognate AUG codon. After start codon recognition, inorganic phosphate (P_i_) is released from the eIF2 complex, which is stimulated by eIF5 acting as a GTPase-activating protein, likely *via* an arginine-finger mechanism (Algire *et al*, 2005; Das *et al*, 2001; Paulin *et al*, 2001). Subsequently, initiation factors apart from eIF1A and eIF3 dissociate (Mohammad *et al*, 2017; Sha *et al*, 2009) and subunit joining with the 60S LSU is then mediated by the GTPase eIF5B.

An important regulatory and scaffolding role in these processes is taken on by the multisubunit complex eIF3 (Cate, 2017; Hinnebusch, 2006), which can be structurally divided into the so-called PCI-MPN core and the more peripheral subunits. In yeast, the PCI-MPN core consists of the two subunits eIF3a (Rpg1/Tif32) and eIF3c (Nip1), whereas in mammals, it is formed by an octamer of eIFs 3a, 3c, 3e, 3f, 3h, 3i, 3k and 3l (Valasek *et al.*, 2017). The peripheral subunits consist of the so-called yeast-like core (YLC) module, containing eIF3b (Prt1), eIF3g (Tif35), and eIF3i (Tif34), as well as the C-terminus of eIF3a, the N-terminal domain of eIF3c that interacts with eIF1 and eIF5 (Valasek *et al*, 2003; Valasek *et al*, 2004; Yamamoto *et al*, 2005; Zeman *et al*, 2019), and in mammals eIF3d. In addition, eIF3j is associated with eIF3 but does not belong to its core, and plays a special role (Block *et al*, 1998; Valasek *et al*, 1999). It was shown that eIF3j participates during termination by recycling eRF3 (Beznoskova *et al*, 2013) and during ribosome recycling by assisting ABCE1 in subunit splitting (Young & Guydosh, 2019). Furthermore, it is involved in dissociation of mRNA from the 40S subunit (Pisarev *et al*, 2007; Pisarev *et al.*, 2010). In the context of initiation, eIF3j is believed to participate in the recruitment of eIF3 to the 40S (Elantak *et al*, 2010; Fraser *et al*, 2004; Nielsen *et al*, 2006), to antagonize premature mRNA recruitment (Fraser *et al*, 2007), and to regulate start-site selection (Elantak *et al.*, 2010).

For a better mechanistic understanding of this complicated interplay, a number of cryo-EM structures of 43S PICs and partial 48S ICs gave first insights into the architectural variety of initiation complexes (Aylett *et al*, 2015; des Georges *et al*, 2015; Eliseev *et al*, 2018; Erzberger *et al*, 2014; Hashem *et al*, 2013; Hussain *et al*, 2014; Llacer *et al*, 2015; Llacer *et al*, 2018; Mancera-Martinez *et al.*, 2017). During 43S assembly, the 40S subunit gets prepared to thread the mRNA into the mRNA binding channel between the 40S body and the head. The main constriction for mRNA is at the so-called “latch”, a structural element formed between ribosomal RNA (rRNA) helix h18 and ribosomal protein (r-protein) uS12 on the 40S body, and h34 and uS3 on the head (Schluenzen *et al*, 2000). Empty or only ABCE1-bound 40S usually don’t adopt a defined head conformation and the latch is rather closed (Heuer *et al.*, 2017; Passmore *et al*, 2007). Binding of eIF1 and especially eIF1A, which bridges the body with the head, seems to prime and confine the 40S by inducing a small rotation of the 40S head (Llacer *et al*, 2015; Passmore *et al.*, 2007), but the latch still remains in a closed position (Llacer *et al.*, 2015). Latch opening was only observed in *in vitro* reconstituted partial 48S ICs containing mRNA and both eIF3 and the eIF2 TC in addition to eIF1 and eIF1A (Llacer *et al.*, 2015; Llacer *et al*, 2018). Here, two conformations of the 48S IC can be distinguished: the open P_OUT_ and the closed P_IN_ conformation, which differ in the orientation of the 40S head and the TC. Compared to the empty and eIF1/1A-bound structures, the head is moved upwards away from the body in the P_OUT_ conformation. This leads to widening of the latch and the P site tRNA_i_ in the TC is only bound via the anticodon loop (AL) to the 40S head but not the body. In the P_IN_ conformation, the AL moves down and engages in stable codon-anticodon interactions with the cognate start codon in the P site, accompanied by a downward movement of the 40S head.

In all eIF3-containing structures, the PCI-MPN core was located on the back of the 40S subunit, from where peripheral subunits stretch out. In 43S PICs, the YLC was found close to the mRNA entry site of the 40S (Aylett *et al*, 2015; des Georges *et al*, 2015; Eliseev *et al*, 2018; Erzberger *et al*, 2014), however only at low resolution. Moreover, the YLC module has been shown to relocate to the intersubunit space (ISS), as observed in *in vitro* reconstituted partial 48S complexes (Llacer *et al.*, 2015), thereby occupying the position of ABCE1. The other peripheral subunits eIF3d and the eIF3c N-terminal domain have been localized near the mRNA exit site (eIF3d: Eliseev *et al.*, 2018) and in the ISS (eIF3c-NTD: Llacer *et al.*, 2015; Obayashi *et al*, 2017). Interestingly, two structures of partial native 43S/48S complexes exist in which ABCE1 could be visualized in substantial quantities (Simonetti *et al*, 2016, re-interpreted in Mancera-Martinez *et al*, 2017; Heuer *et al.*, 2017). Notably, both samples were obtained after adding non-hydrolyzable AMP-PNP and/or GMP-PNP to either yeast (Heuer *et al.*, 2017) or rabbit reticulocyte (Simonetti *et al.*, 2016) lysates and subsequent isolation of the 43S peak from a sucrose gradient. This may have led to non-physiological locking of ABCE1 on the 40S subunit, thereby limiting any conclusions about a putative role of ABCE1 during the phase connecting recycling with initiation. Furthermore, apart from a low-resolution cryo-EM map (Aylett *et al.*, 2015) no structural data exist on eIF3j in the context of the native 43S PIC. Therefore, the native structural landscape enabling the transition from translation termination *via* recycling to initiation is not yet well-understood.

## Results

In this work, we set out to provide a structural inventory of ABCE1-containing 43S or 48S (pre-) initiation complexes from native small ribosomal subunits (SSU). We first asked if substantial amounts of ABCE1 are associated with initiation factor-bound 40S under native conditions. To that end, lysates from a yeast strain (*S.c.)* containing TAP-tagged ABCE1 (Rli1p) were subjected to density gradient centrifugation followed by Western Blotting of fractions (Suppl. Fig. 1A). In agreement with previous studies (Andersen & Leevers, 2007; Pisarev *et al.*, 2010; Pisareva *et al.*, 2011), we observed that ABCE1 was especially enriched on 40S and 80S ribosomes. We further performed affinity purification under varying buffer conditions but without any stabilizing non-hydrolyzable ATP analogs, and analyzed the elution fractions by quantitative mass spectrometry (LC-MS/MS) (Fig. 1A and Suppl. Fig. 1B, 1C). We found that the expected SSU proteins but also eIF3 core components and especially eIF3j (Hcr1), were enriched by ABCE1 affinity purification, indicating that both proteins were indeed integral components of native (pre-)initiation complexes. Because of this finding and since eIF3j was implicated in ABCE1-dependent ribosome splitting *in vivo* (Young & Guydosh, 2019), we tested if eIF3j together with ABCE1 had a direct impact on ribosome splitting in a reconstituted system. To this end, we performed *in vitro* splitting assays in yeast and tested if eIF3j can play a stimulatory role. Purified 80S ribosomes were incubated with the purified splitting factors Dom34, Hbs1, Rli1 (ABCE1), eIF6 to prevent re-association of ribosomal subunits, ATP and GTP as well as different amounts of eIF3j. Splitting efficiency was assessed from sucrose density gradient UV-profiles by monitoring 80S versus ribosomal subunit amounts (Fig. 1B, C and Suppl. Fig. 1D). Indeed, we observed that an addition of eIF3j in molar excess increased the ratio of split subunits when compared to a reaction containing the splitting factors only (Fig. 1C, Suppl. Fig. 1D). Increasing amounts of eIF3j resulted in higher splitting activity. However, eIF3j alone did not exhibit any activity. In addition, we found that eIF3j and substoichiometric amounts of ABCE1 remained bound to the 40S after splitting (Suppl. Fig. 1E). To further confirm that eIF3j can still be associated with the 40S-ABCE1 complex after splitting, we employed the “facilitated splitting” assay as described before (Heuer *et al.*, 2017). In this assay, ribosomes are allowed to dissociate under splitting-promoting conditions (low Mg^2+^ and high salt) and in the presence of putative subunit-binding factors (see Materials and Methods). Indeed, in this assay we observed that eIF3j remained on the 40S SSU together with ABCE1, confirming that the two factors don’t only collaborate during splitting but also remained together on the 40S for downstream events such as initiation (Suppl. Fig. 1F, G).

**Figure 1:**
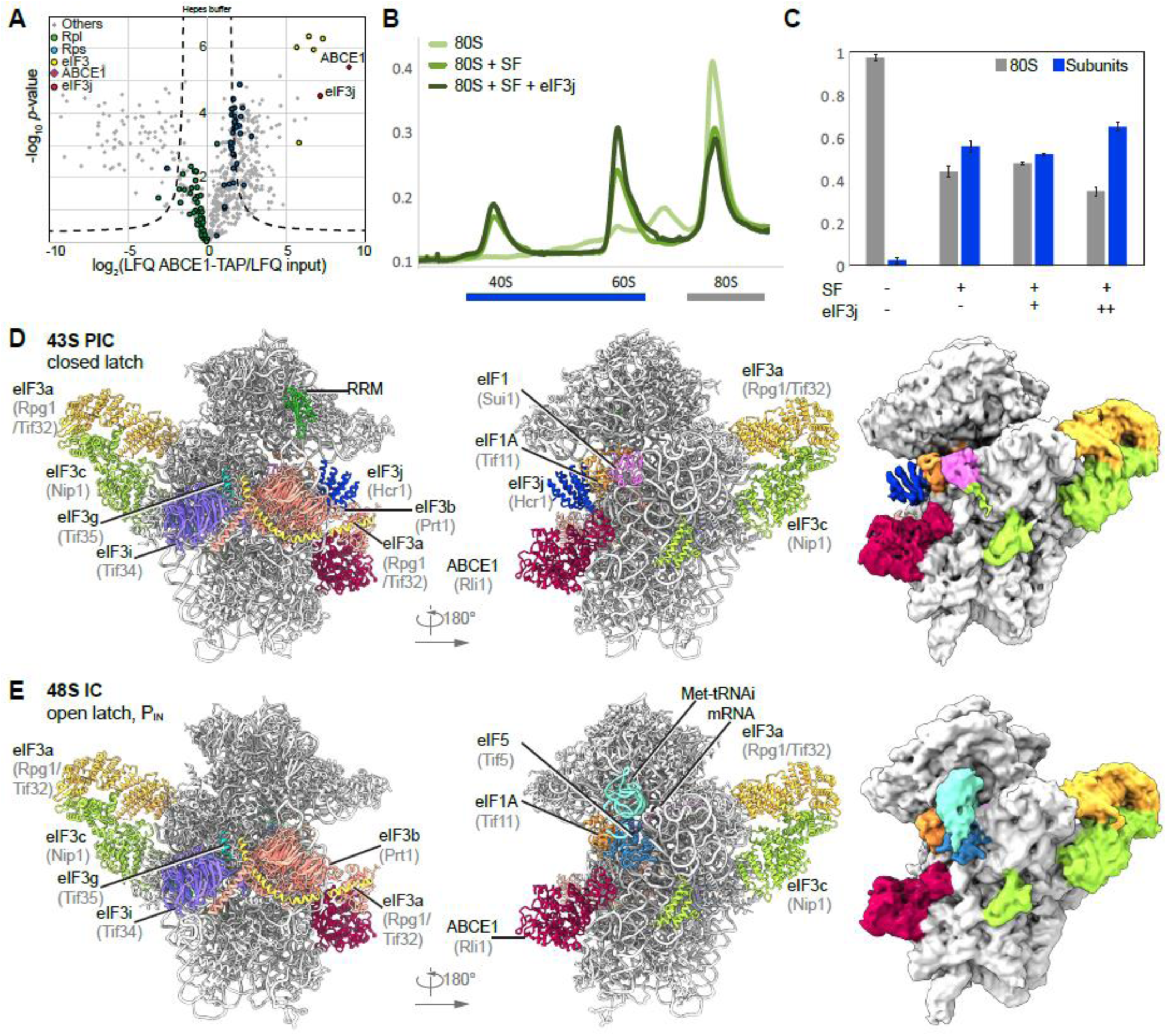
Biochemical analysis and cryo-EM structures of yeast ABCE1-containing initiation complexes. **(A)** Volcano plot representing the statistical analysis of the fold enrichment of proteins after affinity purification in Hepes buffer of ABCE1-TAP followed by label-free quantification (LFQ) using liquid chromatography–tandem mass spectrometry (LC-MS/MS). Proteins above the curved lines show a statistically significant enrichment according to the t test value. **(B)** Sucrose density gradient UV-profile after in vitro splitting assays and **(C)** relative abundance of 80S and subunits as calculated from triplicates; SF = splitting factors including Dom34, Hbs1, ABCE1 and eIF6. **(D, E)** Cryo-EM maps low-pass filtered at 6 Å and models of the yeast subclasses representing an ABCE1- and eIF3j-containing 43S PIC (D) and an ABCE1- and eIF5-containing partial 48S IC (E).

To gain further insights into the composition of native small subunits from yeast and human cell lysates we adopted a shotgun cryo-EM approach. Yeast SSU complexes were obtained after harvesting the crude 43S/48S peak from a preparative sucrose density gradient of yeast cell lysate that was not further treated or stabilized with a non-hydrolyzable nucleotide analog. Similarly, human native 40S were obtained from untreated lysates of HEK Flp-In 293 T-Rex cells after serendipitous non-specific enrichment on sepharose material during unrelated affinity pullouts (see Materials and Methods). Of these samples, large enough cryo-EM data sets were collected in order to analyze their complex composition by extensive 3D classification (Suppl. Fig. 2 and Suppl. Fig. 3).

In the yeast data set, as expected, the selected particles contained (pre-)initiation complexes, which could be further classified into defined states varying in composition and conformation of eIF-associated 40S subunits. The majority of these complexes (62%) contained ABCE1, and the most interesting classes consist of 43S particles containing ABCE1, eIF3, eIF1, eIF1A and eIF3j on the 40S (Aylett *et al.*, 2015; Heuer *et al.*, 2017). The mRNA path (latch) is in the closed conformation (Passmore *et al.*, 2007), and at the mRNA entry we find a density for a typical RNA recognition motif (RRM) (see below). Importantly, in these classes we observed a direct interaction between the FeSD of ABCE1 and eIF3j (Fig. 1D). Moreover, we found one class of particles with mRNA bound, apparently representing a partial 48S IC complex. It contained eIF3, eIF1, tRNA_i_ in the P_IN_ conformation, as well as the N-terminal domain (NTD) of eIF5 as observed before (Llacer *et al.*, 2018), and, to our surprise, also ABCE1 (Fig. 1E). The classes representing 43S PIC and 48S IC were refined to a resolution of 5.3 and 6.2 Å respectively, allowing us to fit molecular models of existing structures as rigid bodies (Fig. 1D, 1E, Suppl. Fig. 4 and Table 1).

**Table 1:** Data collection, refinement, and model composition of the *S.c.* initiation complexes.

|  | **S.c. 43S PIC**<br>**PDB XXXX EMDb XXXX** | **S.c. 48S IC**<br>**PDB XXXX EMDb XXXX** |
| --- | --- | --- |
| **Data collection** |  |  |
| Voltage (kV) |  | 300 |
| Electron exposure (e<sup>-</sup>/Å<sup>2</sup>) |  | 25 |
| Defocus range (μm) |  | -1.1 to -2.3 |
| Pixel size (Å) |  | 1.084 |
| Symmetry imposed |  | C1 |
| **Refinement** |  |  |
| Particle images (no.) | 20,618 | 12,937 |
| Map resolution (Å) | 5.3 | 6.2 |
| FSC threshold |  | 0.143 |
| Map sharpening B factor (Å<sup>2</sup>) | -48.5 | -100 |
| **Model composition** |  |  |
| Correlation coefficient (%; Phenix) | 0.69 | 0.6 |
| Models used (PDB codes) | 5NDG, 6FYY, 4U1E, 3BPJ, 6FEC | 6FYX, 4U1E, 6TB3 |
| Non-hydrogen atoms | 76,429 | 79,327 |
| Protein residues | 8,028 | 7,965 |
| RNA bases | 1,719 | 1,870 |

In the human sample we also found 40S subunits associated with initiation factors, similar to the yeast sample. After classification, four major stable eIF3-containing classes could be obtained (Fig. 2A). The 40S in State I resembled the state of an empty 40S subunit with a closed latch (Heuer *et al.*, 2017; Passmore *et al.*, 2007), and only the core eIF3 subunits and weakly bound eIF1 were found. State II had a similar conformation and we found extra densities in the ISS for eIF1, eIF3j and ABCE1. State III additionally contained eIF1A and the ternary eIF2-GTP-tRNA_i_ complex (TC) in the open P_OUT_ conformation (Llacer *et al.*, 2015), whereas State IV was similar to State III but lacked ABCE1. Notably, in contrast to the yeast sample, we did not find any 48S classes containing mRNA. Thus, our human sample mainly represented 43S post-splitting or pre-initiation complexes prior to mRNA recruitment.

**Figure 2:**
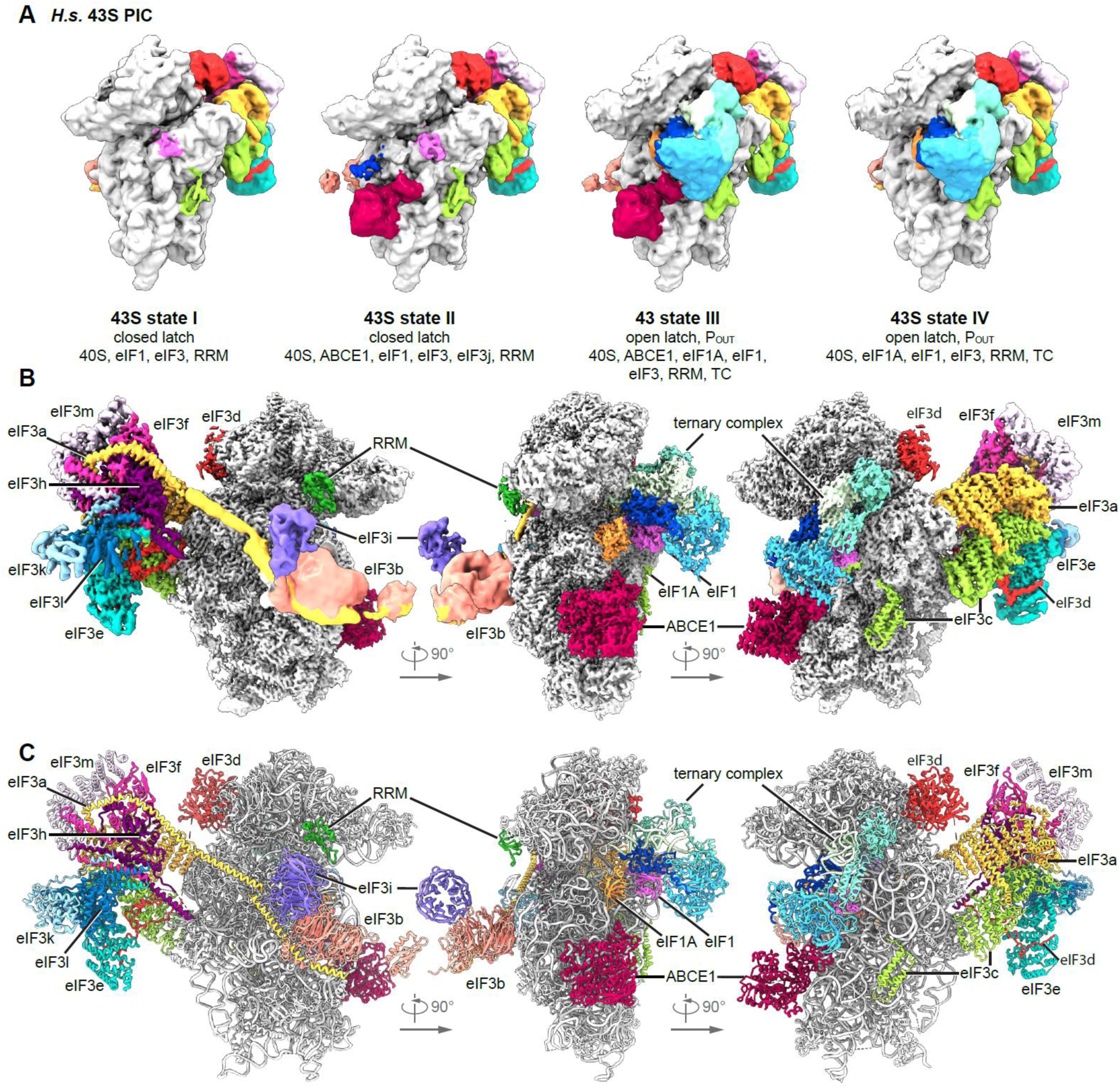
Cryo-EM structures of human 43S PICs in different assembly states. **(A)** Overview of four selected compositional states present in the human 43S PIC data set; **(B)** Composite map of the complete human 43S PIC after focused and multi-body refinements on individual sub-complexes, filtered at local resolution. **(C)** Composite model of the complete human 43S PIC, as represented by state III.

Independent focused classification and multi-body refinements focusing on individual sub-complexes (Suppl. Fig. 3 and Suppl. Fig. 5) enabled us to obtain molecular resolution for large parts of the human 43S sub-complexes. Therefore, we were able to build models for the octameric eIF3 PCI-MPN core at the back side of the 40S, parts of the YLC at the mRNA entry site and most factors located in the ISS, including ABCE1, eIF3j, eIF1 (including the N-terminal tail), eIF1A, the full eIF2 TC, and the eIF3c N-terminal domain, thus resulting in a near-complete molecular model of the human 43S particle bound to ABCE1 (Fig. 2B, 2C, Table 2).

**Table 2:**
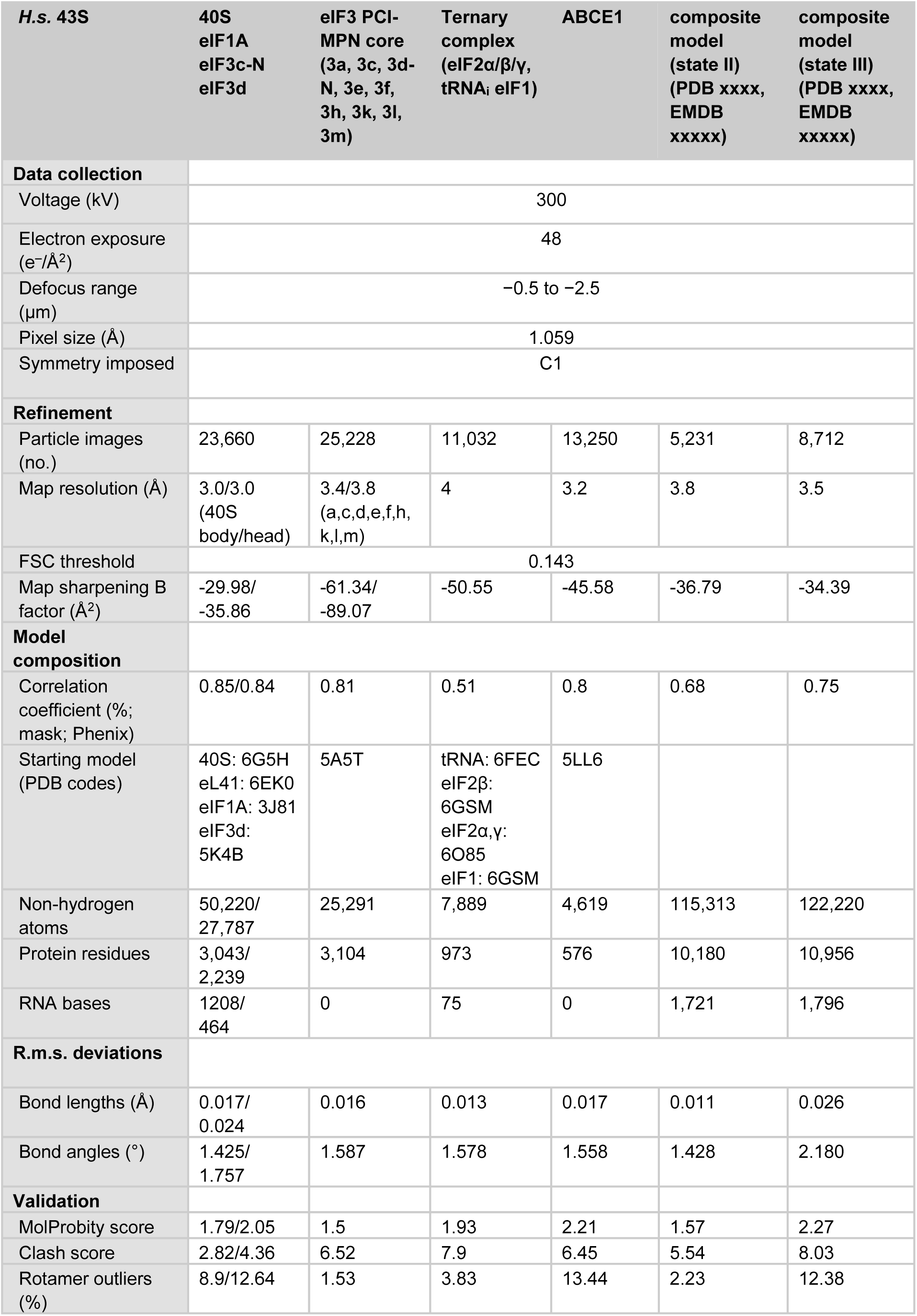
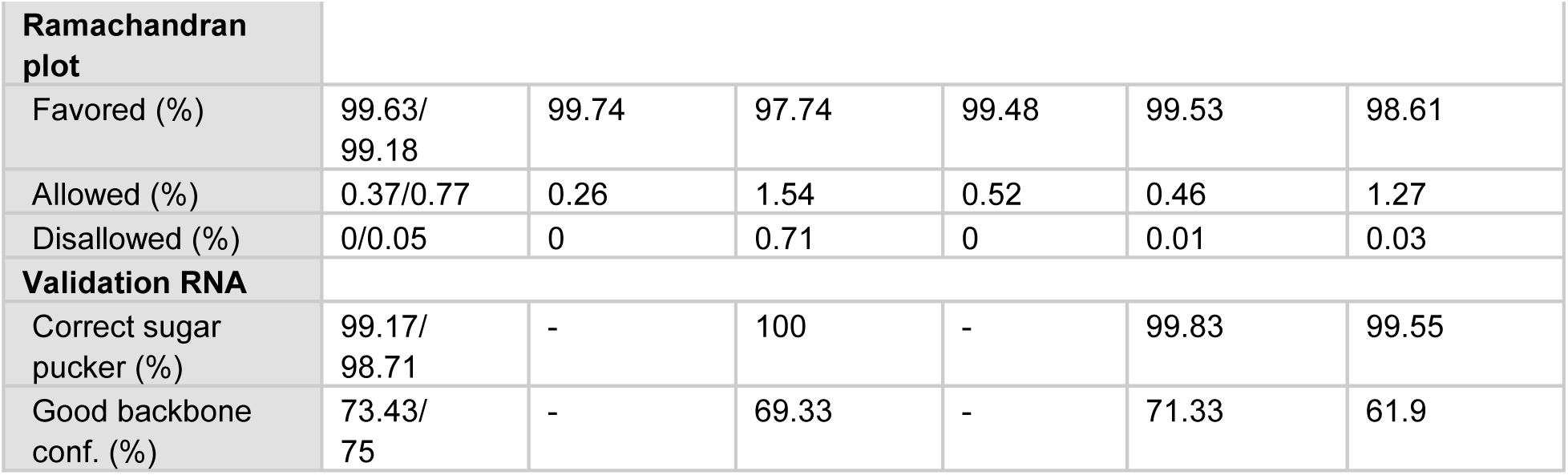
Data collection, refinement, and validation statistics of the human 43S PIC. Atomic models were built into the best-resolved maps as obtained after local focused refinement or multi-body refinement. Validation statistics are shown for each individual part, as well as for the final composite models. The model for State II includes 40S SSU, eIF1, eIF3 PCI-MPN core, eIF3d, eIF3c-N, eIF3a-C, eIF3b, eIF3i, eIF3j, RRM, ABCE1 and the mRNA channel-blocking peptide. The model for State III includes 40S SSU, eIF1, eIF1A, eIF2α/β/γ, tRNA_i_, eIF3 PCI-MPN core, eIF3d, eIF3c-N, eIF3a-C, eIF3b, eIF3i, RRM and ABCE1.

### Conformation of ABCE1-bound 40S-initiation complexes

Strikingly, we observed ABCE1 associated with 40S subunits during all stages of 43S PIC assembly in human and even with 48S IC complexes in the yeast sample. In all complexes, the FeSD of ABCE1 was in the extended conformation packed against h44, and the ATPase body occupied the universal translation factor binding site on the 40S, which is highly similar to previous observations of non-native complexes (h8-h14 junction; h5-h15 junction) (Heuer *et al.*, 2017; Mancera-Martinez *et al.*, 2017; Nürenberg-Goloub *et al.*, 2020) (Fig. 3A). Here, the 40S subunit is engaged in a very similar way as in the archaeal 30S-ABCE1 structure (Nürenberg-Goloub *et al.*, 2020) *via* the ABCE1-specific helix-loop-helix (HLH) domain and the open conformation with respect to the composite hinge region (h1 and h2). Surprisingly, however, in all structures we observed the ATPase in a novel state that has not yet been described for ABC-type ATPases (Fig. 3B-D and Suppl. Fig. 6A): compared to the closed conformation as observed in *in vitro* reconstituted 30S and 40S PSCs (Heuer *et al.*, 2017; Nürenberg-Goloub *et al.*, 2020), we find that only NBSII is closed whereas NBSI adopts a half-open conformation comparable to the one observed in several 80S pre-splitting complexes (Fig. 3B) (Becker *et al.*, 2012; Brown *et al.*, 2015; Preis *et al.*, 2014). When analyzing our best-resolved human map, which was obtained after focused classification on ABCE1, we unambiguously identify an Mg^2+^-ATP (Fig. 3E) occluded in NBSII, similar to the archaeal 30S-ABCE1 structure with Mg^2+^-AMP-PNP (Nürenberg-Goloub *et al.*, 2020). In the human structure, residues of the typical conserved motifs of ABC-type ATPases are involved: Lys386 of the Walker A, Gly220 of the NBD1-Signature loop and His521 of H-loop contact the *γ*-phosphate, and the Mg^2+^ ion is coordinated by Thr387 (Walker A) and Gln415 (Q-loop). In contrast, for NBSI we observed Mg^2+^-ADP bound exactly as observed in the crystal structures of open archaeal ABCE1 (Barthelme *et al.*, 2011; Karcher *et al*, 2008): Y87 of the A-loop stacks on the adenine base, F92 on the ribose, the Walker A loop (Asn112-Ser117) binds the *α*- and *β*-phosphates, and the Mg^2+^ ion is coordinated by the *β*-phosphate, Ser117, Gln171 (Q-loop) and Asp241, Glu242 (Walker B). Importantly, the Signature loop of NBD2 (Leu463-Glu467), which occludes ATP in the catalytically active closed state, is moved by 3.5 Å away from NBD1. In conclusion, our data suggest that - in contrast to the nucleotide-occluded state observed *in vitro* - in native SSU-ABCE1 complexes, ATP-hydrolysis in NBSI has already occurred, whereas NBSII is still inhibited.

**Figure 3:**
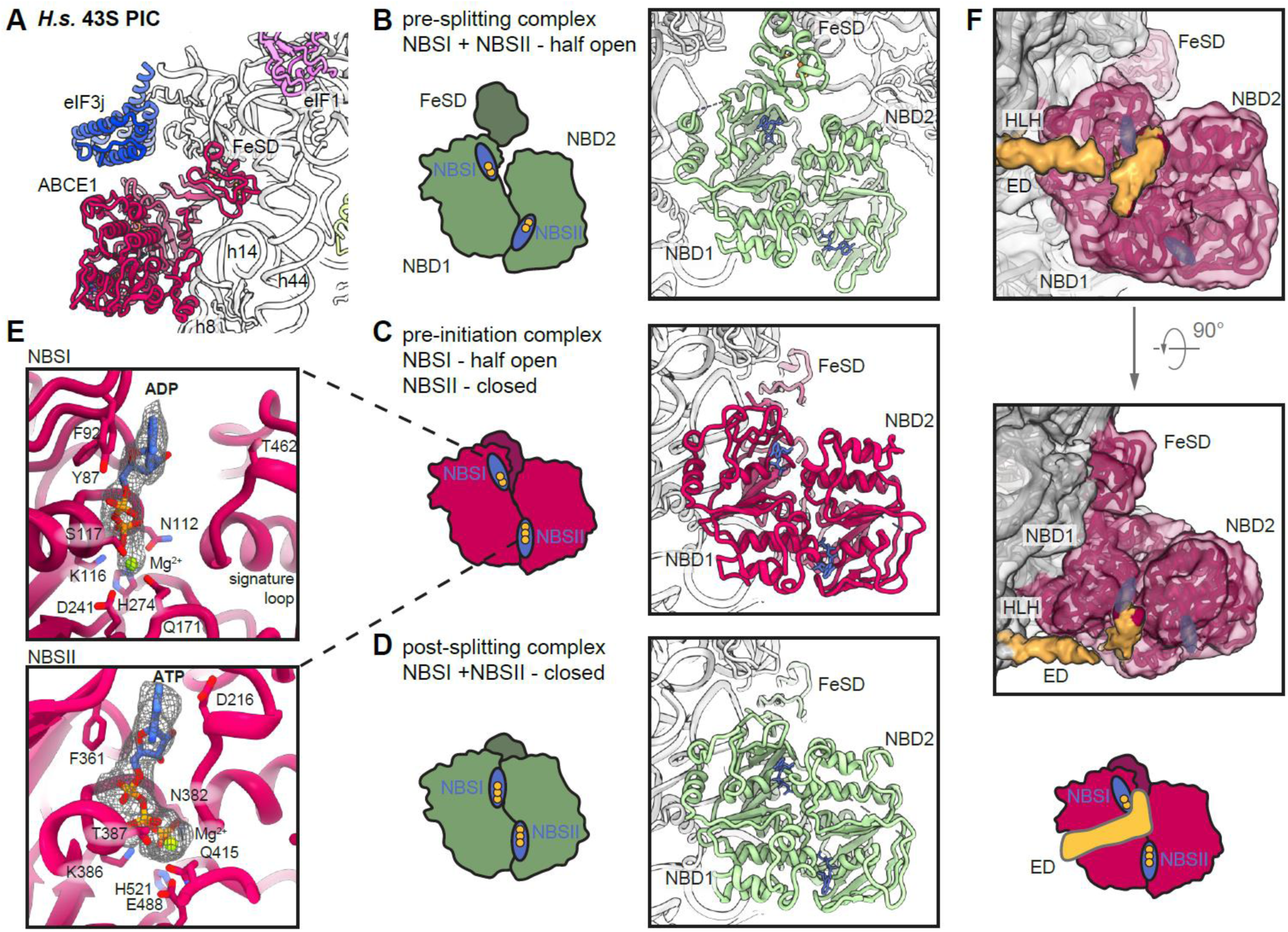
Conformation of ABCE1 in native 40S initiation complexes. **(A)** Overall position of ABCE1 in 40S initiation complexes, here representatively shown for the human State II with eIF3j. **(B-D)** Schematic representation and structure of semi-open ABCE1 as in 80S-pre-splitting complexes (Brown et al., 2015, PDB 3JAH) (B), hybrid semi-open/closed ABCE1 as in native 40S initiation complexes (C) and fully closed ABCE1 as in in vitro reconstituted post-splitting complexes (Nürenberg-Goloub et al., 2020, PDB 6TMF) (D). **(E)** Zoom into NBSI and NBSII showing bound Mg2+-ADP (in NBSI) and Mg2+-ATP (in NBSII) fit in density as obtained after focused classification on ABCE1 and refinement. **(F)** View focusing on the NBDs and the unassigned extra density (ED) reaching from the 40S via the HLH into NBSI. The ABCE1 map was low-pass filtered at 6 Å. Schematic representation highlighting the position of the ED with respect to the NBSs.

As an additional difference to previous structures, we observed a rod-like extra density (ED) after low-pass filtering in all native 43S PIC structures, protruding from h17 of the 40S body into the cleft between NBD1 and NBD2 of ABCE1 (Fig. 3F). In the human complex, it contacted the HLH motif and residues C-terminal of the following Q-loop of NBD1, as well as residues adjacent to the Signature loop of NBD2. Unfortunately, we were not able to assign the identity; however, this unknown factor is in a position that easily allows for modulation of the ATPase activity of ABCE1 by restricting further movements of the HLH or the two NBDs with respect to each other. Interestingly, the position of the ED on ABCE1 is similar to the one observed in a recent structure of archaeal ABCE1 co-crystallized with an 18-mer fragment from the C-terminus of the archaeal 60S stalk protein aP1 (Imai et al., NAR 2018). This would be consistent with an idea that ABCE1 may possess a multivalent interaction patch in this region, which would allow for regulation of its ATPase activity. In the post-splitting situation, one plausible candidate for contributing the density between the NBDs is eIF3j, which was previously shown to interact with ABCE1 (Fraser *et al.*, 2007; Khoshnevis *et al*, 2010; Kispal *et al*, 2005). Moreover, the eIF3j subunit appears to already work in concert with ABCE1 during ribosome recycling, both *in vivo* (Young & Guydosh, 2019) and - as shown above (Fig. 1B, 1C) - *in vitro.* Regardless of the identity of the factor, the observed stabilization of ABCE1 in the half-open conformation with one non-hydrolyzed ATP still bound may indeed indicate an inhibition of ATPase activity, which would explain its rather stable association with the 40S subunit.

Consistent with this idea and with biochemical data, we found yeast and human 43S PIC sub-populations concomitantly bound to ABCE1 and eIF3j. The eIF3j subunit was positioned on the intersubunit side, roughly resembling the location previously described in low-resolution maps (Aylett *et al.*, 2015) (Fig. 4). The main difference between the maps was the absence (human) or presence (yeast) of eIF1A. However, apart from a small rotation around the neck (approx. 3°) we did not observe significant conformational changes of the 40S when comparing the two structures. In both structures, we could fit the crystal structure (or a homology model for yeast, Suppl. Fig 6B, 6C, 6D) of the human eIF3j dimer (unpublished; PDB 3BPJ; lacking 137 residues at the N- and 28 residues at the C-terminus) essentially without adjustments (Fig. 4B, 4D). In brief, the dimer folds into a stable entangled 6-helix bundle that is arranged such that the N-termini are in close vicinity but C-termini face into opposite directions.

**Figure 4:**
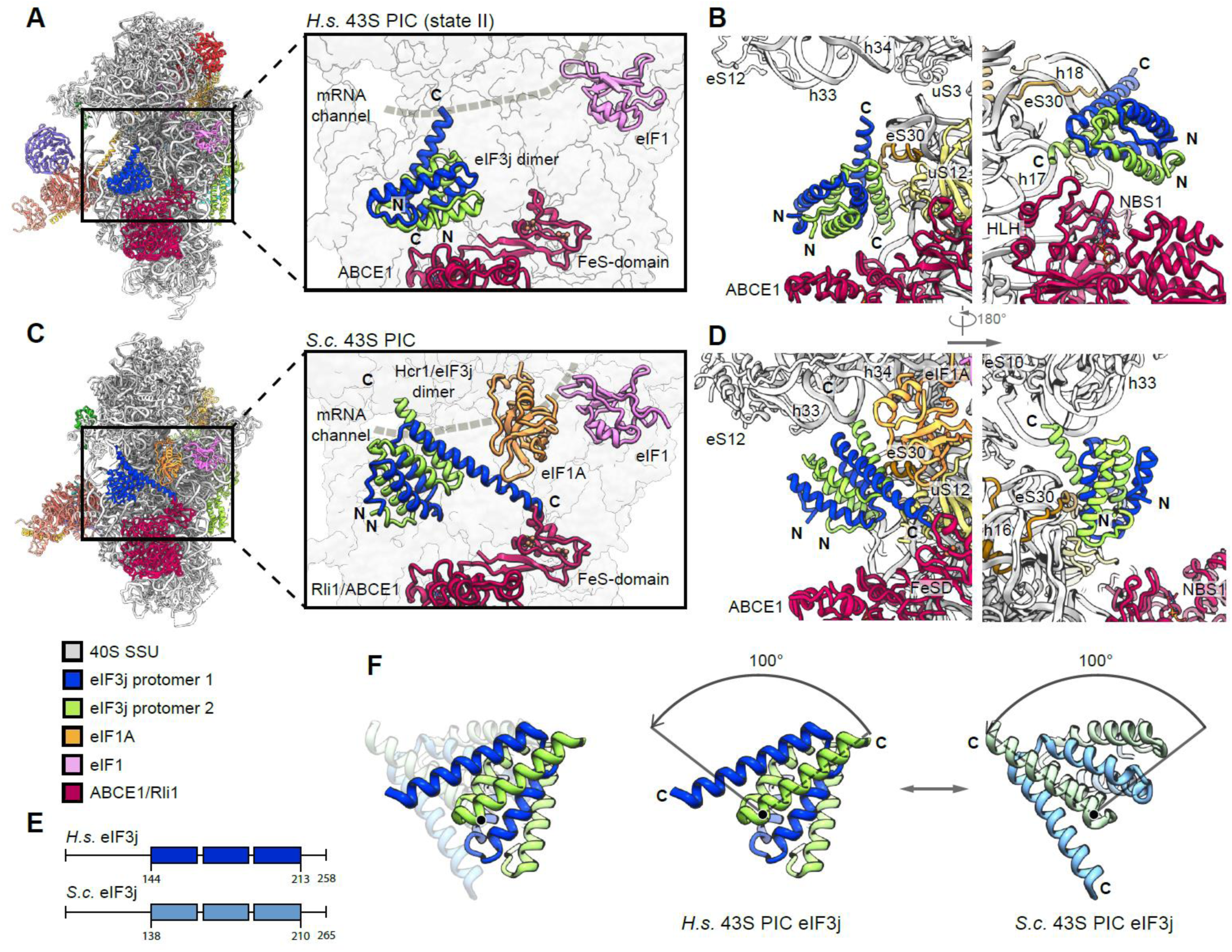
Two conformations of eIF3j in human and yeast 43S PICs. **(A)** Overview and zoomed view on the model of human 43S PIC II (lacking eIF1A), focusing on the two protomers of the dimeric eIF3j 6-helix bundle in the ISS. eIF3j is in close vicinity to NDB1 of ABCE1 but only forms contacts to the 40S. The mRNA channel is indicated by a dashed gray line. **(B)** Two different views showing the interaction of the two *Homo sapiens* (*H.s.*) eIF3j protomers with the 40S. **(C)** Same views as in (A) on the model of the yeast 43S PIC. Here, eIF3j (Hcr1) is turned approximately 100 degrees around a pivot formed by the C-terminal helices contacting eS30 and uS12. Protomer 1 thereby contacts eIF1A and the FeSD of ABCE1 and protomer 2 contacts h33. **(D)** Two different views showing the interaction of the two *S.c.* eIF3j protomers with the 40S and ABCE1. **(E)** Schemes of *H.s.* and *S.c.* eIF3j indicating the length of N- and C-termini; the three helices present in the crystal structure of human eIF3j (PDB 3BPJ) are indicated. **(F)** View highlighting the rearrangement (100-degree rotation) of eIF3 in the human (eIF1A-lacking) and yeast (eIF1A-containing) 43S PICs.

In the low-pass filtered human State II, which lacks eIF1A, we identified the 6-helix bundle located above the ABCE1 ATPase body and in close vicinity to NBD1 (Fig. 4A, 4B), but no direct contacts were formed with ABCE1. On the 40S, eIF3j contacted the N-terminal tail of eS30 (protomer 1) and the C-terminus of uS12 (protomer 2). The C-terminal helix of protomer 2 further projects towards the three-way junction formed by h32, h33 and h34 at the 40S head, whereas in protomer 1 it points towards h17 and the HLH of ABCE1 (Fig. 4B and Suppl. Fig. 6C). In this position, the N-termini of eIF3j are located above the ABCE1 ATPase body close to the NBD1-NBD2 cleft.

In the yeast 43S PIC, in which eIF1A was present, we found eIF3j in a similar position, but different conformation compared to the human structure (Fig. 4C, 4D). Here, the 6-helix bundle is stably anchored between the 40S beak at rRNA h33 on one side and the 40S body near the ABCE1 FeSD and eIF1A on the other side. The two sides of the anchor are formed again by the C-terminal helices of eIF3j: protomer 2 contacts eS30 at a similar site as in the human structure (Ser16-Thr18) but now the entire helix bundle was rotated by approximately 100 degrees (Fig. 4E, 4F). Consequently, the tip of the protomer 2 C-terminal helix now pointed towards the 40S head, where it contacted the minor groove of rRNA h33 (Fig. 4D and Suppl. Fig. 6D). The C-terminal helix of protomer 1 projected towards the ABCE1-FeSD. First, it passed along eIF1A (likely contacting the loop Glu30-Gln33) and extended further to contact the FeSD *via* Met34 and/or Gly35 by extending the helical secondary structure by 2 turns (from Arg215-Thr221 when starting with the crystal structure). Additional contacts to the 40S were formed by the N-terminal helix 1 of protomer 1 (to the h17-h18 junction; A542; A544) and the loops between protomer 2 helix 1 and helix 2 (to uS12) as well as helix 2 and helix3 (to h33; U1262). At the same time, the N-termini pointed towards the solvent side. Notably, eIF3j would not directly clash with the mRNA path in either of the observed conformations. However, its proposed role in antagonizing mRNA binding *in vitro* (Fraser 2007) may be explained by a stabilization of the closed-latch conformation of the early 43S PIC.

Taken together, the comparison between these two structures suggests that ABCE1 and eIF3j may act in conjunction with eIF1A during key steps of translation initiation.

### Molecular architecture of the PCI-MPN core and its interactions with 40S

State I of the human sample represented a stable class with mainly eIF3 and weak density for eIF1 bound to the 40S SSU. This appears plausible when considering that eIF3 activity during termination and ribosome recycling has been proposed (Beznoskova *et al*, 2013; Pisarev *et al.*, 2007; Valasek *et al.*, 2017), which further indicates that eIF3 can already bind the 40S before eIF1A comes into play. The lack of ABCE1 in this complex may be a result of fast dissociation after splitting or of an alternative splitting mechanism. In any case, after accommodation of eIF1 and eIF1A, the eIF2 TC binds to the 43S to induce the P_OUT_ conformation (State III-IV). Here, the improved resolution allowed us to describe the interaction network of these factors at unprecedented molecular detail.

The PCI-MPN core is located at the back-side of the 40S as observed before (des Georges *et al.*, 2015; Hashem *et al*, 2013; Srivastava *et al*, 1992), and high resolution of the core was obtained by multi-body refinement of State I and State II particles. The structure assembles into β-sheets with the shape of an arc formed by PCI domains of subunits eIF3a, c, e, l, k and m. The arc wraps around a seven-helix bundle formed by the C-terminal helices of subunits c, e, f, h, k, and l (Fig. 5A and Suppl. Fig. 7D), resulting in the typical five-lobed structure (left and right arm, left and right leg and head), which was visualized at a local resolution of 3.4 Å (left arm, head and right arm) and 3.8 Å (left and right leg) (Suppl. Fig. 5C). This allowed for an almost complete molecular interpretation (Suppl. Fig. 7D and Table 3), thus refining previous low-resolution models (des Georges *et al.*, 2015; Eliseev *et al.*, 2018; Erzberger *et al.*, 2014), for example by correcting the register of helices and extending molecular models.

**Figure 5:**
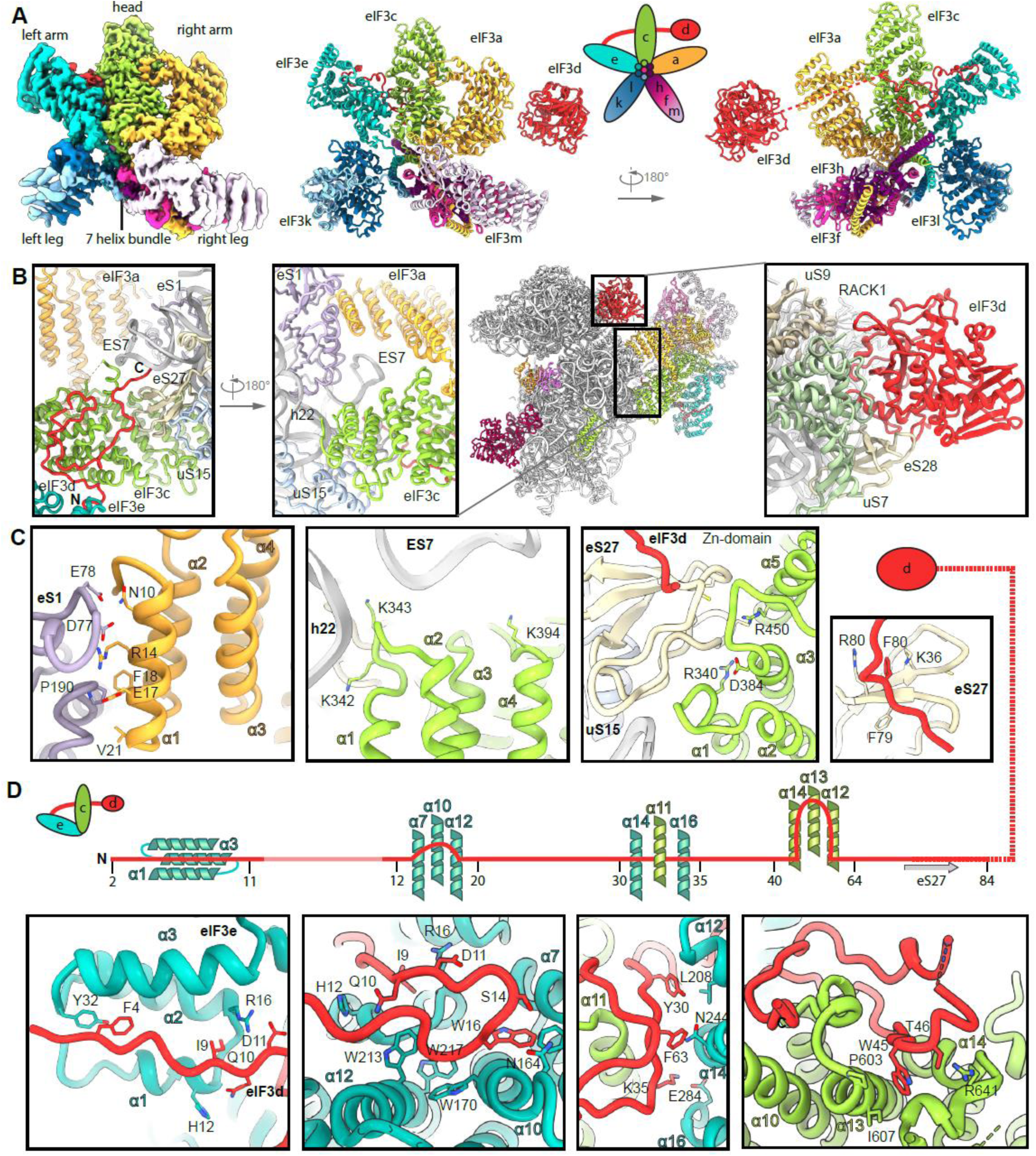
Molecular interactions of the human PCI-MPN core of eIF3 in the 43S PIC. **(A)** Isolated map and molecular model of the eIF3 PCI-MPN core color coded as in Fig 2. Structural hallmarks are indicated, and a scheme shows the composition of the lobes. **(B, C)** Interactions of eIF3a, eIF3c and eIF3d with the ribosome: (B) shows an overview of the structure and zoomed views highlighting the interactions of eIF3a, eIF3c, the eIF3d N-terminal tail and the eIF3d cap-binding domain with the 40S, (C) shows molecular details of eIF3a interacting with eS1; eIF3c interacting with rRNA h22 and eIF3c and the N-terminal tail of eIF3d with the Zn-knuckle domain of eS27. **(D)** Interactions of the eIF3d N-terminal tail with the PCI-MPN core.

**Table 3:** Molecular interactions between eIF3 subunits and 40S.

| eIF3a |  |  |  |  |  |  |  |  |  |
| --- | --- | --- | --- | --- | --- | --- | --- | --- | --- |
| 3a | eS1 | 3a | 3c | 3a | 3f | 3a | 3h | 3a | 3m |
| Q6 | E78 | E125 | K474 | N521 | S232 | N521 | S232 | S444 | K342 |
| N10 | E78 | V132 | N678 | H565 | R109 | A525 | Q314 | I446 | I325 |
| R14 | E78, D77 | R140 | P464, S466 |  |  | E547 | K220 | Y447 | K342 |
| E17 | N75 | R143 | D463 |  |  | H550 | W216 | Q448 | I325 |
|  |  | W246 | P729 |  |  | E564 | H209 | S449 | D326, Q327 |
|  |  | Q247 | Q724 |  |  | R571 | E77 | E451 | Q327 |
|  |  | D337 | K745 |  |  | E576 | R108 | R454 | E309 |
|  |  | L342 | R719 |  |  | R578 | E145, E146 | N512 | C323 |
|  |  | E468 | Y799 |  |  |  |  |  |  |
|  |  | R483 | S797, D800 |  |  |  |  |  |  |
|  |  | I484 | D800 |  |  |  |  |  |  |

| eIF3b |  |  |  |
| --- | --- | --- | --- |
| 3b | 18S rRNA | 3b | uS4 |
| K487 | A560 | R507 | D158 |
| D488 | A560 | V556 | K155 |

| 3c-N |  | eIF3c (PCI-MPN core) |  |  |  |  |  |  |  |  |  |  |  |
| --- | --- | --- | --- | --- | --- | --- | --- | --- | --- | --- | --- | --- | --- |
| 3c | 18S rRNA | 3c | 18S rRNA | 3c | eS27 | 3c | 3d | 3c | 3e | 3c | 3f | 3c | 3h |
| R47 | C1039, A1181 | K342 | G929 | G341 | Q65 | D562 | A70 | Y583 | Y286 | Q852 | D333 | Y557 | L210 |
| K55 | C1180 | K343 | G929 | N388 | Q75 | M591 | Q31 | D587 | Y286 | N870 | N351, Q347 | L859 | M247 |
| K92 | U1178 |  |  | L389 | Q75 | H593 | P32, G64 | D602 | Y41 |  |  | K862 | M247 |
| K132 | C369 |  |  |  |  | Q595 | L39 | S820 | N302 |  |  | V873 | N261 |
| K136 | U367 |  |  |  |  | H600 | H73 | L835 | I347, Q349 |  |  | H876 | N261 |
| Q143 | U367, U368 |  |  |  |  | D602 | H73 | Q837 | S352 |  |  |  |  |
| K144 | U367 |  |  |  |  | P603 | W45 | H845 | N395 |  |  |  |  |
|  |  |  |  |  |  | Q606 | W45 |  |  |  |  |  |  |
|  |  |  |  |  |  | Y609 | K41 |  |  |  |  |  |  |
|  |  |  |  |  |  | N610 | A43 |  |  |  |  |  |  |
|  |  |  |  |  |  | N631 | R38 |  |  |  |  |  |  |
|  |  |  |  |  |  | D635 | Y59 |  |  |  |  |  |  |
|  |  |  |  |  |  | R641 | T46 |  |  |  |  |  |  |
|  |  |  |  |  |  | E666 | W45 |  |  |  |  |  |  |

| eIF3d |  |  |  |  |  |  |  |  |  |
| --- | --- | --- | --- | --- | --- | --- | --- | --- | --- |
| 3d | 3e | 3d | uS27 | 3d | 18S rRNA | 3d | eS28 | 3d | uS7 |
| F4 | Y32 | E75 | K36 | R212 | C1470 | Q416 | R51, E52 | K472 | Q29 |
| Q10 | H12 | E77 | R80 |  |  | T423 | R13 | S478 | N31 |
| D11 | R16 | Q81 | S78, F79 |  |  | K426 | F34, T38 |  |  |
| W16 | S167, G171 | V83 | G76 |  |  |  |  |  |  |
| G17 | W170 |  |  |  |  |  |  |  |  |
| A20 | Q209 |  |  |  |  |  |  |  |  |
| Y30 | L208 |  |  |  |  |  |  |  |  |
| F33 | N244, L590 |  |  |  |  |  |  |  |  |
| K35 | Q283, E284 |  |  |  |  |  |  |  |  |

| eIF3e |  |
| --- | --- |
| 3e | 3l |
| R369 | D480 |
| Y401 | I523 |
| Q416 | D532 |

| eIF3f |  |  |  |  |  |  |  |
| --- | --- | --- | --- | --- | --- | --- | --- |
| 3f | 3h | 3f | 3k | 3f | 3m | 3f | 3l |
| R108 | D113 | S267 | Q218 | Q280 | N362 | N332 | I523 |
| V163 | Y99 | N328 | E203 | D301 | W347 |  |  |
| S164 | E56 |  |  | R306 | H339 |  |  |
| Y239 | L218, E217 |  |  | N313 | S337 |  |  |
| N260 | K206 |  |  |  |  |  |  |
| R261 | H209, N207 |  |  |  |  |  |  |
| I263 | I205 |  |  |  |  |  |  |
| D268 | Q336 |  |  |  |  |  |  |
| Q271 | S161 |  |  |  |  |  |  |
| P316 | Q348 |  |  |  |  |  |  |
| I318 | Q348 |  |  |  |  |  |  |
| N342 | K331 |  |  |  |  |  |  |
| Q345 | N324 |  |  |  |  |  |  |
| N356 | R313 |  |  |  |  |  |  |

| eIF3h |  |  |  |
| --- | --- | --- | --- |
| 3h | 3l | 3h | 3k |
| E195 | R545 | K331 | S217 |
|  |  | K345 | K204 |

| eIF3l |  |
| --- | --- |
| 3l | 3k |
| K534 | S216 |

The main anchor of the eIF3 PCI-MPN core to the 40S is provided by the eIF3a and eIF3c subunits, which form the “head” and the “right arm” of the PCI-MPN core, respectively. eIF3a contacts eS1 *via* its N-terminal PCI-helix H1 and the loop between H1 and H2. Here, Arg14 forms salt-bridges to Glu78 and Asp77 of eS1 (see Table 3 for an inventory of observed molecular interactions). A second contact site was established between Glu17, Phe18 and Val21 of eIF3a and the eS1 Pro190 as well as adjacent residues. The loop H1-H2 of eIF3c (residues 340-345) interacts with rRNA h22 (G929, C930) and multiple sites at the Zn-knuckle domain of eS27 (Fig. 5C and Suppl. Fig. 7C). Furthermore, the *β*-sheet insert between PCI helix 4 and 5 (residues 417-441) of eIF3c forms interactions with uS15, and basic residues in the PCI loops of both eIF3a and eIF3c are positioned to interact with the flexible tip of rRNA ES7 (Fig. 5B).

An additional anchor of the eIF3 PCI-MPN to the 40S is provided by the N-terminus of eIF3d (from A2 to D84) (Fig. 5C, 5D and Suppl. Fig. 7A). Interestingly, we found that it meanders along the PCI helices 1 to 3, 7, 9, 10 and 12 of eIF3e (left arm) and bridges eIF3e with eIF3c (head) by interacting with PCI helices 12, 14 and 16 (eIF3e) and PCI helix 11 (eIF3c). Another specific contact between eIF3c and eIF3e is formed by stacking of Y286 (eIF3e) to Y583 (eIF3c). Moreover, eIF3d also interacts with PCI helix 10, 13 and 14 of eIF3c by forming a large loop, which is anchored by the conserved Trp45 (interactions to Pro603, Ile607 and Glu666 of eIF3c). The interaction to eS27 is established *via* its Zn knuckle, where Phe80 of eIF3d is sandwiched between the side chains R80 and K36 of eS27.

Taken together, the PCI-MPN core of eIF3 establishes a multi-modal molecular interaction pattern with the 40S involving the eIF3a, c and d subunits, which display an unexpected degree of inter-connectivity.

### Structure and location of the peripheral subunits

The peripheral subunits, which consist of the YLC, the eIF3c-NTD, and in humans the eIF3d cap-binding protein domain, are connected to the PCI-MPN scaffold *via* flexible linkers. While eIF3a connects *via* its CTD to the YLC module located close to the mRNA entry site, the N-terminus of eIF3c protrudes from the mRNA exit towards the ISS, where it interacts directly with eIF1. While the N-terminus of eIF3d as an integral part of the PCI-MPN core is anchored to the 40S body, the cap-binding protein domain of eIF3d is located on the 40S head close to the mRNA exit site as observed before (Eliseev *et al.*, 2018). Here, it contacts the 40S SSU *via* its highly conserved helix α10 (Lee *et al*, 2016) that packs upon eS28 *via* Gln416, Thr423 and Lys426 and reaches into the interface between eS28 and uS7, where Gln416 stacks on Arg51 (eS28), which in turn stacks on Phe61 (uS7). The eIF3d helix α12 lies on top of uS7 and forms contacts *via* Lys472, Glu475, Ser478, and Gln479. Notably, since eIF3d is bridging the 40S head with the eIF3 PCI-MPN core anchored to the 40S body, it could serve to relay conformational rearrangements of the 40S head - as occurring during the assembly of 43S and 48S complexes - to the PCI-MPN core or, vice versa, allow the eIF3 complex to directly control the conformational state of the 40S head. (Fig. 5D and Suppl. Fig. 7A, 7B).

For eIF3c, only a part of its NTD could be located on the ISS of the 40S so far, where it forms a helix bundle (Llacer *et al.*, 2015). We found a particularly stable arrangement of the eIF3c NTD in classes containing the eIF2 TC and, after multi-body refinement, local resolution of 3 to 4 Å (Suppl. Fig. 5B and Suppl. Fig. 8A, 8B) allowed us to determine the register of the four eIF3c-NTD helices (Val47 to Y149) (Fig 6). A stretch preceding the first helix (47-51) contacts h24 and h27 *via* R47 to the backbone phosphate of C1039 and the 2’-OH of A1181. The peptide bond of Val49 of eIF3c stacks on base C1180, which is also contacted by the first helix (52-74) of the bundle. Here, the two charged residues K55 and R56 interact with the backbone of rRNA G1179 and C1180. Backbone-phosphate interactions were also formed by the second helix (76-92) to rRNA h11 (A364) and h27 (U1178), by the fourth helix (136-143) to rRNA h11 *via* K136 (to U367), and finally by the peptide bond of Thr140 stacking upon the U367 base, as well as Gln143 hydrogen-bonding to U367. Additional but less rigid contacts were established by the K-rich loop between helix 3 and helix 4 of eIF3c (Fig. 6E, 6F, Suppl. Fig. 8A, 8B and Table 3).

**Figure 6:**
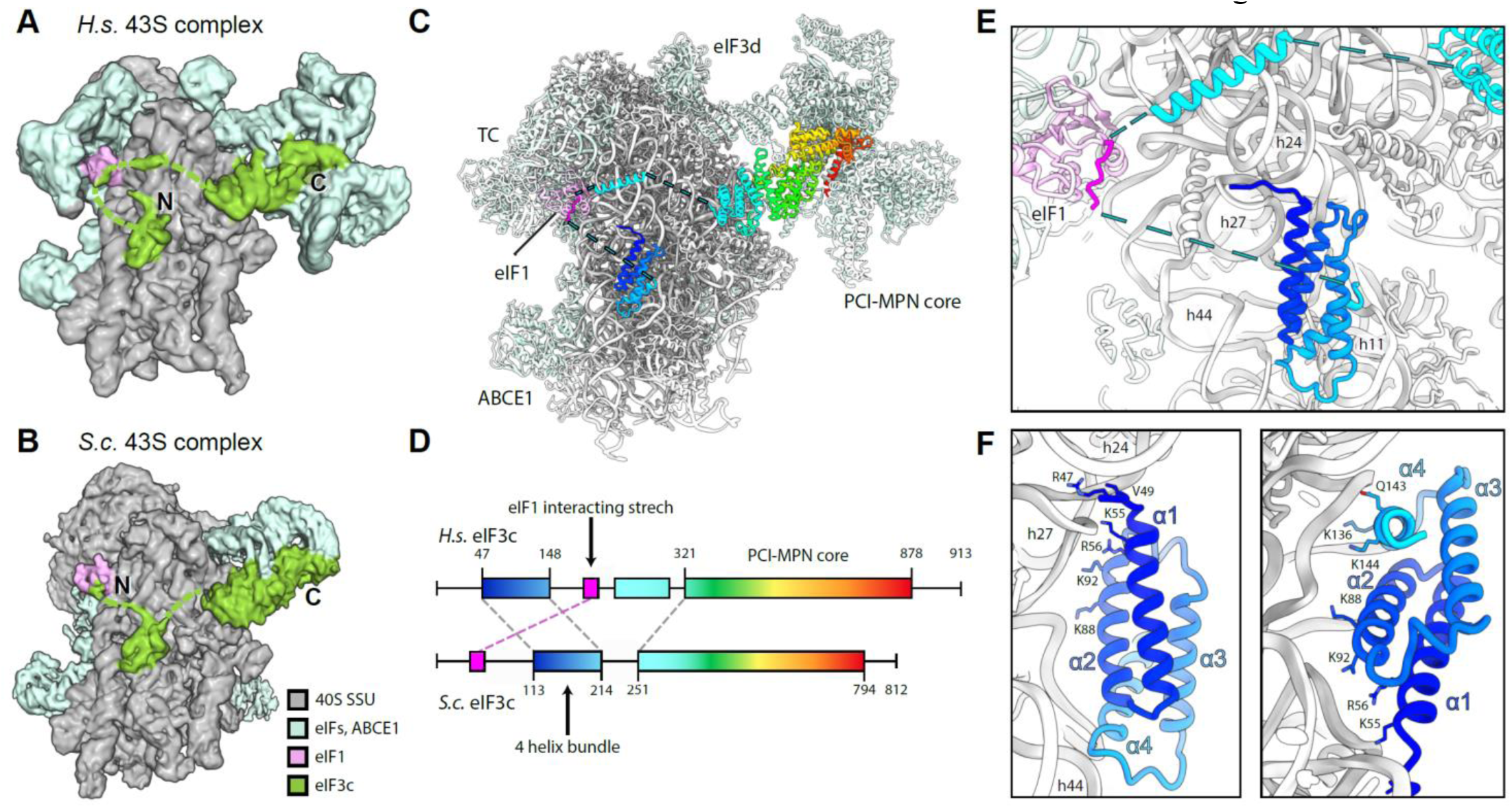
Arrangement of the eIF3c-NTD in human and yeast 43S PICs. **(A)** Cryo-EM map obtained after focused sorting of the human 43S PIC on the TC: when low-pass filtered at 6 Å, it shows the density of almost-complete eIF3c-NTD in the ISS. **(B)** Cryo-EM map of the yeast 43S PIC low-pass filtered at 6Å. **(C, D)** Model for human eIF3c in the TC-containing 43S colored in rainbow (C) and scheme of the alignment between human and yeast eIF3c sequences, colored accordingly (D). The eIF1-interacting stretch present in the N-terminus of *S.c.* eIF3c shows 32.0/56.0% sequence identity/similarity with an insert C-terminal of the conserved 4-helix bundle conserved in mammals. **(E)** Zoomed view highlighting the position of the eIF3c NTD and eIF1 in the 40S ISS. **(F)** Molecular model for the 4-helix bundle interacting with 40S rRNA and r-proteins.

Notably, when low-pass filtered, a rod-like extra density for the eIF3c N-terminus became apparent, bridging the 4-helix bundle with eIF1 near rRNA h23 and h24. This density was neither present in our nor in other (Llacer *et al.*, 2015) yeast 43S/48S reconstructions, where the four-helix bundle was directly connected to the eIF3c core moiety, and a site N-terminal of this region interacted with eIF1 (Fig. 6A, 6B). Sequence alignments of the yeast and human eIF3c N-termini revealed an insertion on the C-terminal side of the conserved four-helix bundle in human (Fig. 6C, 6D, Suppl. Fig. 8C). This insertion from residue 165 to 213 displays 32.0% sequence identity and 56.0% sequence similarity with a stretch at the N-terminus of yeast (42-92), which was previously shown to be involved in the interaction of eIF3c with eIF1 by NMR studies (Obayashi *et al.*, 2017). Here, chemical shift perturbation after eIF1 binding is observed for Glu51, Ala67 and a stretch between Lys68 and Lys77. Moreover, in our human complex one stretch of well-resolved density for the eIF3c-NTD was present at the eIF1 loop between helix *α*1 and helix *α*2 (Asp53-Lys58) as well as Ile100 and Gly101 of *α*2 (Suppl. Fig. 8D). This observation is highly consistent with the NMR study, in which the same interacting region on eIF1 is identified for the eIF3c-NTD of yeast. Together, these observations lead us to the conclusion that the density observed near eIF1 in the human structure corresponds to this insertion C-terminal of the helix bundle, fulfilling an analogous role to the previously characterized N-terminal stretch of eIF3c in yeast.

From local classification, we also obtained one class with strong density for the YLC module including the eIF3a-linker that connects it to the PCI-MPN core (Suppl. Fig. 9A). In brief, the YLC module contains two *β*-propellers: the 7-bladed WD40 repeat of eIF3i and the 9-bladed WD40 repeat near the C-terminus of eIF3b. The two propellers are held together by the C-terminal helical domain of eIF3b, which is formed by 3 *α* - helices: the most C-terminal one binds to eIF3i, while the two preceding *α*-helices are bracketing the eIF3a C-terminus against the eIF3b *β*-propeller (des Georges *et al.*, 2015; Herrmannova *et al*, 2012). N-terminal of its *β*-propeller, eIF3b contains a noncanonical RNA recognition motif (RRM) (ElAntak *et al*, 2007) that can form further interactions with the eIF3a-CTD (Dong *et al*, 2013; Khoshnevis *et al*, 2014; Valasek *et al*, 2002; Valasek *et al*, 2001) as well as the N-terminus of eIF3j (Elantak *et al.*, 2010; Valasek *et al.*, 2001).

For the CTD of eIF3a we could build a long *α*-helix (residues 602-743) into the elongated rod-like density protruding from the PCI-MPN core to contact uS2 and eS21 (Suppl. Fig. 9A). This helix extends further towards the YLC where it forms a hinge-like structure and then connects to the stretch of the eIF3a helix that is bound to the eIF3b *β*-propeller. It thereby contacts the tip of the otherwise flexible rRNA expansion segment ES6C, which in turn contacts the loop between the first two helices of the eIF3b helical domain. In this arrangement, the eIF3b WD40 is rigidly confined between rRNA h16 and uS4 on one side, and ES6C on the other side, and is thus well resolved in the proximity of the 40S (Suppl. Fig. 9B, Table 3). The eIF3i-eIF3g complex as well as the eIF3b-RRM, however, remained rather flexible as observed before (Erzberger *et al.*, 2014). Nonetheless, we observe a stabilization of the eIF3b-RRM in ABCE1- and eIF3j-containing classes, possibly due to an interaction of the eIF3b-RRM with the eIF3j N-terminus (Elantak *et al.*, 2010; Valasek *et al.*, 2001).

In yeast, the positioning of the YLC module at the mRNA exit was the same, because here it was also held in place by ES6C (Suppl. Fig. 9C). However, in the majority of particles in the yeast dataset (approximately 85%), we could observe a conformational change of the eIF3i-eIF3g module relative to the ES6 anchor point. Especially in the eIF3j-containing 43S class, the eIF3i-eIF3g entity rotates by approximately 120 degrees away from the mRNA entry towards ES6C and ES6B. The loop preceding the eIF3i-contacting helix of eIF3b (Thr697-Asp701) appears to serve as a hinge for this rotation (Suppl. Fig. 9D).

Apart from the YLC, we observed an additional density near the mRNA entry at the tip of h16 in all of our 43S structures, which was previously assigned to the RRM of eIF4B (Eliseev *et al.*, 2018) (Suppl. Fig. 10A). This density is especially prominent in subclasses of the human dataset lacking the TC, in which we could unambiguously identify the typical RRM fold at a local resolution around 4 Å (Suppl. Fig. 10B, 10C). Notably, besides eIF4B, the largely flexible eIF3g subunit is a potential candidate for this density because it also contains an RRM, which shares very high structural and sequence similarity (50.0%) to eIF4B (Suppl. Fig. 10D 10E), and it was cross-linked to the nearby proteins uS10 and uS3 (Cuchalova *et al*, 2010). Unfortunately, at the current resolution we cannot unambiguously distinguish these two RRMs in our maps and it is possible that both compete for the same binding site. Next to this domain, we observed density reaching from the RRM into the mRNA channel in all early 43S PIC structures with a closed latch (Suppl. Fig. 10A 10B). Close to the RRM, this density forms a loop that shows multiple contacts to uS3 before winding along uS3 towards the mRNA channel. Within the channel, one side chain can clearly be identified as a tryptophan facing towards uS3 (contacting Lys148 and Met150) and further interacting with uS3 Leu142 and Val115. The stretch also contacts 18S rRNA G626, A628 and U630 of h18 as well as C1698 of h28, C1331 and A1489 of h34 (all in the A site). Thereby, this peptide stretch blocks the entire mRNA channel down to the P site where it contacts the flipped-out base C1701 at the tip of h44. Unfortunately, local resolution in this region is insufficient to provide further molecular detail and clearly identify this entity. However, it is apparent that accommodation of mRNA in the 48S IC complex would require its relocation, which may allow for allosteric communication between the different eIFs.

### Conformation of the ternary complex

After analyzing the eIF3 complex, we also gained molecular information on the human eIF2 TC by focused classification. The TC as well as eIF1 and eIF1A were observed on the intersubunit side in a similar overall position and conformation as described before for other PICs in P_OUT_ conformation at low resolution (PDB 6GSM, PDB 3JAQ (Llacer *et al.*, 2015)) (Suppl. Fig. 3). Briefly, eIF2 consists of three subunits, *α, β* and *γ*. The eIF2*γ* subunit shares structural homology to EF-Tu-like translational GTPases (e.g. Schmitt, EMPB J 2002) and consists of a G-domain (domain I), including the regulatory switch loops (swI and swII), followed by two *β*-barrel domains. eIF2*α* consists of an N-terminal OB-fold domain, a central helical domain, and a C-terminal α-β domain. The eIF2*β* subunit has an unstructured N-terminal domain, followed by a central helix-turn-helix (HTH) domain and C-terminal zinc binding domain (ZBD). In solution, tRNA_i_ was shown to be bound to the TC in a distinct way different to canonical tRNA-bound EF-Tu/eEF1A by employing additional composite interactions with both eIF2*α* and eIF2*γ* (Schmitt *et al*, 2012). The eIF2*β* subunit, however, has never been sufficiently resolved to elucidate its molecular contribution to tRNAi binding and 43S PIC formation.

In our structure, we found the tRNAi embraced by all three eIF2 subunits (Fig. 7A, 7B). Similar to the 5 Å resolution crystal structure (3V11 Schmitt et al. 2012), the methionylated CCA end is sandwiched between the GTPase domain and domain II of eIF2*γ*. The terminal adenine base A76 is accommodated in a pocket formed by the *β*-sheets of the eIF2*γ* domain II including Val278, Phe322, Gly340 and Arg260 (Fig. 7C, Suppl. Fig. 8C). The 2’-OH group of the ribose moiety interacts with the carbonyl group of Ala323 and the methionyl side chain stacks on Tyr83 of eIF2*γ* G domain. The CCA-end is further stabilized by contacts including a cation-*π* stack of Lys266 on tRNA_i_ C75 and Asn71 of the eIF2*γ* Sw1 loop with tRNA_i_ C74. Moreover, Arg296 of the eIF2*β* ZBD intercalates into the major groove of the acceptor-stem helix (G70; supported by Lys293 contacting the phosphate backbone of U69) (Fig. 7D, Suppl. Fig. 8C). eIF2*α*-contacts the T- and D-loops mainly *via* its central helical domain whereas the N-terminal OB-fold domain intercalates between anticodon stem and uS7 in the E site on the head of the 40S. The central eIF2*β*-HTH domain contacts the anticodon from the A site and thereby forms multiple contacts to eIF1, also involving residues of the newly built C-terminus (I314-R329), which stretches below the tRNA_i_ anticodon stem towards the E site and contacts C1057 of rRNA h24 (*via* N327).

**Figure 7:**
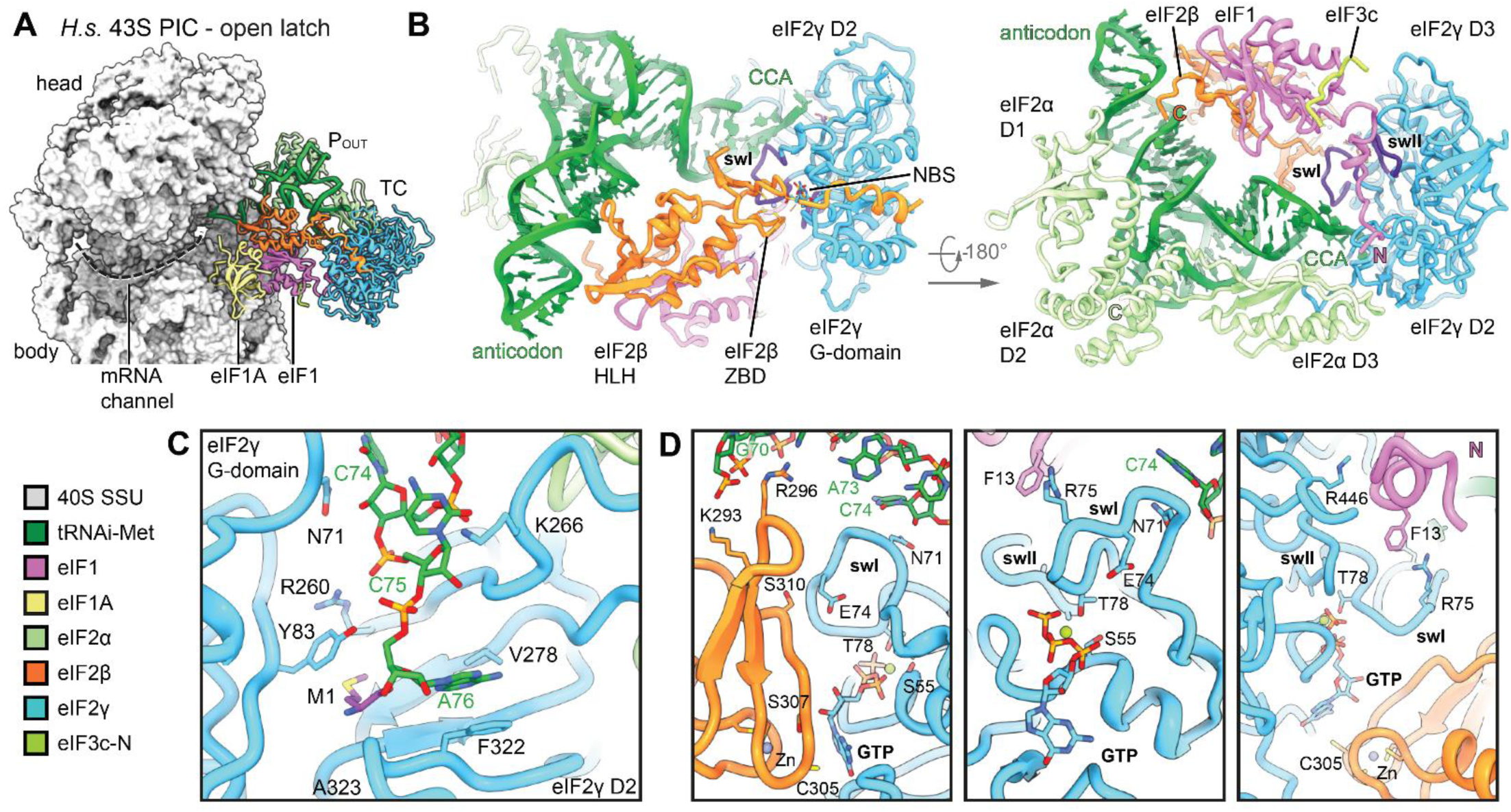
Conformation of the TC in the complete human 43S PIC. **(A)** Overview highlighting the positions of TC, eIF1 and eIF1A in the complete human 43S PIC. **(B)** Interactions of eIF2 subunits and domains and eIF1 with methionylated tRNAi; switch loops (sw) of eIF2γ are labeled and colored in purple; the *de novo* built N-terminal tail of eIF1 and the C-terminus of eIF2α and eIF2β are labeled with N and C, respectively. **(C)** Molecular interactions of the methioninylated CCA-end of tRNAi and eIF2γ. **(D)** Molecular interactions within the nucleotide binding pocket and conformation of sw loops stabilized by the eIF1 N-terminal tail, the eIF2β ZBD and tRNAi.

Notably, in the GTP binding pocket of eIF2*γ* we clearly identified a Mg^2+^-GTP (Fig. 7D). Ser55 of the conserved P-loop and Thr78 of swI coordinate the Mg^2+^-ion, whereas Asp134 and Pro135 of swII likely contact the *γ*-phosphate. Compared to the crystal structure of the archaeal TC (Schmitt et al., 2002), the guanine base is rotated by 90° and accommodated in a pocket between Asn190 and Ala226 of eIF2*γ* and Cys305 of the eIF2*β*-ZBD, which is tightly packed upon the nucleotide binding pocket.

Interestingly, both switch loops were embedded in a tight interaction network involving interactions with tRNA_i_, eIF2*β* and the eIF1 N-terminal tail, which we built *de novo.* The N-terminal tail of eIF1 protrudes from the 5-stranded *β*-sheet and binds to Arg446 of eIF2*γ* domain III, where it forms a loop and projects towards Arg75 of eIF2*γ* swI, forming a cation-*π* stack with Phe13 (Fig. 7D and Suppl. Fig. 8C, 11D). Furthermore, the conformation of the swI loop was stabilized by the tRNA_i_ *via* Asn71 (see above) and an interaction between conserved Ser310 of the ZBD of eIF2*β* with Glu74.

In close vicinity to the guanosine binding pocket, we find eIF2*β* Ser307, the equivalent of yeast eIF2*β* Ser264. In yeast, a Ser264Tyr mutation causes the Sui^-^ (suppressor of initiation codon) phenotype, leading to increased utilization of UUG start codons (Huang *et al*, 1997). This mutation was shown to increase GTP hydrolysis rates and stabilize the closed P_IN_ conformation of the 43S PIC (Martin-Marcos *et al*, 2014). In the observed position, the tyrosine mutation of Ser307 could easily interfere with the bound nucleotide, for example by stacking on the guanine base, and thus alter the geometry of the nucleotide binding pocket.

Taken together, we found the TC in a stable state within the 43S PIC, in an open conformation in absence of mRNA. An intricate interaction framework is established by the 40S and eIF1 to accommodate the GTP-bound eIF2-tRNA_i_ in a rigid position. The switch loops are kept in a rigid conformation stabilized by tRNA_i,_ eIF2*β* and the eIF1 N-terminal tail, and the GTPase pocket of eIF2*γ* is closed by eIF2*β*. This may prevent premature release of the bound nucleotide and, at the same time, may restrict access for eIF5-NTD to avoid premature GAP activity.

Following TC assembly on 43S PIC and opening of the latch, mRNA can be threaded into the mRNA binding site, followed by scanning for the first AUG codon by the 48S particle. While we do not find scanning intermediates in either yeast or human datasets, in our yeast native 40S population we find one state containing eIF1A, tRNA_i_ in the P_IN_ state and the eIF5-NTD instead of eIF1 (yeast 43S PIC). Apart from weaker density for eIF2, this state is similar to one observed before (Llacer *et al.*, 2018), where it was interpreted as a late state after start-codon recognition. However, to our surprise we still find ABCE1 in this complex. This suggests that ABCE1 may play further roles even in later stages of initiation, or that its dissociation is not required at this stage.

## Discussion

While the role of highly conserved ABCE1 during ribosome recycling has been studied in mechanistic details (Becker *et al.*, 2012; Nurenberg-Goloub *et al.*, 2018; Nürenberg-Goloub *et al.*, 2020), its role after 60S dissociation remained largely elusive. However, when first characterized biochemically, ABCE1 was found associated with 43S/48S (pre-)initiation complexes in yeast, humans and *Drosophila* (Andersen & Leevers, 2007; Chen *et al.*, 2006; Dong *et al.*, 2004). Since then, it is a long-standing question what the function of ABCE1 in these complexes is. Our extensive single particle analysis of native small subunits from yeast and human cells shows that ABCE1 can stay associated with the 40S throughout the assembly of the 43S PIC prior to mRNA loading. Surprisingly, in yeast we even find ABCE1-48S complexes beyond the stage of mRNA engagement and start codon recognition as indicated by the presence of the eIF5-NTD.

We further observe that in all ABCE1-containing 43S structures its NBDs are in an unusual hybrid conformation, where NBS2 is closed and NBS1 is semi-open. This is contrary to previous *in vitro* studies showing SSU-associated ABCE1 in the ATP-occluded fully closed state. Notably, the two NBSs in ABCE1 were shown to be highly asymmetric and NBSII has a low ATP-turnover rate compared to NBSI (Gouridis *et al.*, 2019; Nurenberg-Goloub *et al.*, 2018). Consistent with this behavior, we find Mg^2+^-ATP still bound in the closed NBSII, whereas Mg^2+^-ADP is present in NBSI. This is in agreement with the most recent model for the ABCE1 ATPase cycle, in which closure of the NBSII was discussed to be the decisive step for disassembly of 80S pre-splitting complexes, a process that is then triggered by subsequent closure and ATP-hydrolysis in NBS1. Subsequently, re-opening of NBSI would be expected on the small subunit. But if ATP-hydrolysis is prevented either by usage of a non-hydrolyzable ATP analog or by hydrolysis-deficient Walker B mutants, ABCE1 can be trapped in the fully closed state on the small subunit under facilitated splitting conditions (Heuer *et al.*, 2017; Kiosze-Becker *et al.*, 2016; Nürenberg-Goloub *et al.*, 2020). In native ABCE1-associated complexes, however, NBSI is already in a more open conformation and additionally obstructed by a yet unassigned peptide which intercalates between the two NBDs close to NBSI. This factor either keeps NBSI from closing (after putative binding of another ATP), or alternatively, prevents further opening into a state as observed in free ABCE1. This brings up the question of why ATP-hydrolysis in NBSII, which would lead to dissociation from the 40S SSU, is inhibited. We find NBSII in a very similar conformation as in the fully closed archaeal structure (Nürenberg-Goloub *et al.*, 2020), and the structure reveals no clues to explain why ATP-hydrolysis is slowed down. Thus, we speculate that a further and likely only small-scale allosteric signal into NBSII may be necessary for its activation. This may - after dissociation of the obstructing factor - occur upon further opening of NBSI and be accompanied by changes in the ABCE1-specific HLH and hinge regions.

The observation that ABCE1 dissociation can apparently be actively prevented points towards a direct role in 43S PIC and even 48S IC assembly, most likely in concert with eIF3j. We could corroborate the finding that eIF3j assists in ABCE1-dependent splitting by *in vitro* dissociation assays and furthermore we established that eIF3j remains bound to the 40S together with ABCE1 after the splitting cycle. Here, our structures of early 43S PICs suggest that eIF3j and ABCE1 may be beneficial for binding of eIF1A. In the yeast conformation, eIF3j appears like a molecular ruler reading out the exact distance between the post-splitting-specific FeSD conformation of ABCE1 and the 40S head and beak conformation as adopted after eIF1A binding. Thus, it is tempting to speculate that the observed conformational change of eIF3j may play a role in priming the 40S for eIF1A binding and/or stabilizing the early closed-latch conformation of the 43S PIC when eIF1A is bound. Notably, eIF1A is the only factor that was not found to be pre-assembled in a 40S-free multi-factor complex (MFC) consisting of eIF1, eIF2-tRNA_i_-GTP, eIF3 and eIF5 in yeast (Asano *et al*, 2000; Zeman *et al.*, 2019), plants and mammals (Sokabe *et al*, 2012). While eIF1A is capable of binding 40S SSU independently and adopting a similar conformation as within the context of initiation (Yu *et al*, 2009), it is possible that after binding of the MFC a conformational or positional rearrangement may be required for its productive integration into the 43S complex. Concluding our cryo-EM analysis of native initiation complexes, we can deduce a putative order of events during 43S PIC and 48S IC assembly by formation of several structural hallmarks. As a first step the MFC binds to 40S as indicated by the highly populated eIF3-eIF1 bound classes. While the PCI-MPN core is stably anchored at the solvent side of the 40S, the eIF3c-NTD locates into the ISS *via* the 4-helix bundle, positioning eIF1 in the process. The YLC module is guided to the mRNA-entry by stable positioning of the eIF3b *β* -propeller between h16 and of rRNA expansion segment ES6c. Here, the eIF3i-eIF3g complex can adopt variable positions that may be important for the role of eIF3g-eIF3i during scanning (Cuchalova *et al.*, 2010). Concomitantly, the RRM of either eIF3g or eIF4B accommodates on the mRNA entry. We note, that in most classes of early closed latch 43S PICs, the mRNA channel is additionally blocked by an extra density that may be further missing parts of eIF3g, eIF4B, the CTD of eIF3a, or the ribosome hibernation factor SERBP1 (Stm1 in yeast) (Anger *et al*, 2013; Ben-Shem *et al*, 2011; Brown *et al*, 2018). After eIF1A accommodation, the TC can be stably integrated to form the complete mRNA-free P_OUT_ state 43S. This opens up the latch and leads to clearance of the mRNA path, since in P_OUT_ complexes no density in the mRNA path is visible.

With respect to a fully accommodated TC, our structure reveals for the first time a network of interactions between the tRNA_i_ and all subunits of eIF2 as well as eIF1 at molecular resolution. The eIF2*γ* switch loops are highly confined and the GTPase pocket is closed by the ZBD of eIF2*β*, thus restricting the access for the eIF5-NTD to exert its GAP activity. Notably, GTP-hydrolysis in eIF2*γ* may already occur during scanning. This would require that the eIF5 N-terminal tail could reach into the eIF2*γ* GTPase pocket and, thus, result in a rearrangement of the eIF2*β*-ZBD. A structure of a scanning 48S, however, is still lacking. Yet, large structural rearrangements have been observed after start-codon recognition, during which the 48S IC adopts the closed P_IN_ state. Here, the entire TC rearranges, and especially eIF2*β* alters its location on the 40S head and relative to eIF1 and eIF1A. It is likely that this conformational switch could already partially occur during scanning and that this would also affect the position of the eIF2*β*-ZBD, which was too flexible to be resolved in all previous cryo-EM structures (Llacer *et al.*, 2015; Llacer *et al*, 2018; Simonetti *et al*, 2016; Eliseev *et al.*, 2018). After eIF5-dependent GTP-hydrolysis, release of inorganic phosphate (P_i_) would still be inhibited until start codon recognition. During or after this process the eIF5 NTD replaces the gatekeeper eIF1 and leads to a further stabilization and compaction of the P_IN_ state, which may be a prerequisite for the following step of eIF5B-mediated subunit joining (Llacer *et al.*, 2018).

Our analysis shows that ABCE1 can still be associated with initiating 40S. Yet, which role might ABCE1 play during formation of the full 43S and - as observed in yeast - even in context of the eIF5-accommodated partial 48S? Currently, ABCE1 is assumed to act as an anti-association factor, ensuring that premature 60S interaction is prevented after termination and ribosome splitting. However, in this function it would likely become redundant after the formation of the 43S PIC, failing to explain its presence in later stages of initiation. Another possibility is that its observed interplay with eIF3j as early as during the splitting reaction supports the timely recruitment of the remaining eIFs to the vacant 40S. Furthermore, we speculate that the inhibiting peptide close to NBSI would need to be ejected to facilitate ATP-hydrolysis in NBSII. Here it is possible that dynamics of the rather flexible YLC module could play a role. In fact, this module is able to relocate into the ISS to occupy the position of ABCE1 (Llacer *et al.*, 2015). With this steric competition in mind, it would be plausible that it contributes to ABCE1 dissociation, although it is not entirely clear at which stage this relocation happens. In addition, eIF3j, which is still present at least as fuzzy density in the fully assembled 43S, may also contribute in coordinating such events, for example *via* its known interaction with eIF1A and the eIF3b-RRM (Elantak *et al.*, 2010). Finally, since ABCE1 is even present on 48S IC complexes after start codon recognition, events during subunit joining may be the final trigger for ABCE1 dissociation. In this context, the P proteins of the 60S subunit may not only play a role during ribosome splitting as suggested before (Imai *et al*, 2018), but also for ABCE1 removal after initiation. Yet to reveal exact timing of these events and the mechanistic interplay of these factors, future work will be needed.

## Acknowledgements

The authors thank H. Sieber, J. Musial, C. Ungewickell and S. Rieder for technical assistance, L. Kater and K. Best for support with the pre-processing pipeline of cryo-EM data, R. Buschauer for assistance in model building, L.Valášek and A. Jacquier for critical reading of the manuscript and J. Wells for support during the setup of splitting assays.

This work was supported by German Research Council (BE1814/15-1 and TRR174), the Center for Integrated Protein Science Munich (CiPS-M), the ANR-17-CE11-0049-01 and the ANR-17-CE12-0024-02 grants, the Pasteur Institute and the Centre National de la Recherche Scientifique. H.K. is supported by a DFG fellowship through the Graduate School of Quantitative Bioscience Munich (QBM).

## Author contributions

H.K., T. M.-K., T.B. and R.B. designed the study. M.A. prepared the sample for the human and T.M.-K. for the yeast native 40S complexes. T.M.-K. and H.K. prepared components for *in vitro* splitting assays and T.M.-K. performed splitting assays. M.A., T.M.-K. and O.B. collected and M.A. and T.M.-K. processed the cryo-EM data. H.K., J.C. and T.M.-K. built and refined the model. H.K., T.M.-K., T.B. and R.B. analyzed and interpreted the structures. E.D, A.N. and M.F.-R. performed ABCE1-TAP purification and quantitative label-free MS. T.B., H.K., T.M.-K. and R.B. wrote the manuscript with contributions from all authors.

## Materials and Methods

### Yeast Strains

*Saccharomyces cerevisiae* ribosomes for biochemical assays were purified from a wild-type BY4741 strain, which was grown on YPD medium.

Samples for LC-MS/MS analyses were purified from a BY4741 (*MATa, ura3*Δ*0, his3*Δ*1, leu2*Δ*0, met15*Δ*0*), Rli1-TAP:HIS3MX6 strain (Ghaemmaghami *et al*, 2003).

### ABCE1-TAP polysome profile and sucrose density gradient fractionation

Yeast (*Saccharomyces cerevisiae*; *S.c.*) cells from the BY4741 strain expressing C-terminally TAP-tagged ABCE1 (Rli1) were grown in 200 mL YPD to an OD_600_ of 0.8. The cells were then treated with 50 µg ml^−1^ cycloheximide on ice for 5 min. and collected by centrifugation. The cells were lysed in lysis buffer (20 mM Tris-HCl, pH 7.4, 50 mM KCl, 10 mM MgCl_2_, 50 µg ml^−1^ cycloheximide and EDTA-free protease inhibitors (Roche)) by vortexing them with glass beads (12 cycles of 30 sec. vortex/30 sec. on ice). The lysate was cleared by centrifugation for 10 min. at 16,000 × *g*, 4 °C and stored at −80 °C. Ten OD_260_ units were loaded on a 10–50% sucrose gradient and centrifuged at 187,813 × *g* for 2.75 h at 4 °C in a SW41Ti rotor (Beckman Coulter). The fractions of the gradient were collected, and proteins were precipitated with trichloroacetic acid and separated on a 10% acrylamide gel. The proteins were detected with antibodies after western blotting: ABCE1-TAP with peroxidase anti-peroxidase (PAP) complex (Sigma-Aldrich) at 1:2,000, and Nog1 with a rabbit anti-Nog1 antibody at 1:5,000 dilution.

### ABCE1-TAP Tandem Affinity Purifications

Cells expressing C-terminally TAP-tagged ABCE1 were cultivated in rich medium (YPD) until OD_600_ of 2, and cultures were centrifuged at 4 °C, rinsed in cold water, and frozen at −80 °C. Cells were thawed on ice, resuspended in lysis buffer (50 mM Tris/HCl pH 8.0, 100 mM NaCl, 10 mM MgCl_2_, complete EDTA-free protease inhibitor mix or: 20 mM HEPES/KOAc pH 7.4, 100 mM KOAc, 10 mM MgCl_2_, complete EDTA-free protease inhibitor mix), and lysed with glass beads using a Magnalyser. The lysates were clarified by centrifugation at 16,000 × *g* for 10 min. at 4 °C. Supernatants were collected and Triton (0.5% final) or NP40 (0.1% final) was added to the lysate. Binding to magnetic beads coupled with IgG was performed on a wheel at 4 °C overnight. Beads were collected on a magnet, flow-through was discarded and beads were washed in lysis buffer. Elution was performed by resuspension in 2% SDS, 1× Tris-EDTA buffer and incubation at 65 °C for 10 min. Eluted beads were discarded on a magnet and eluate was purified on HiPPR Detergent Removal Resin (Thermo Scientific, 88305). Purified proteins were eluted in PBS. The rest of the eluates was precipitated by the methanol/chloroform technique (Wessel & Flugge, 1984) and analyzed by mass spectrometry.

To control the quality of the affinity purification, a sample of eluates (3%) was separated on acrylamide NuPAGE Novex 4–12% Bis-Tris gels (Life Technologies) and analyzed by silver staining.

### Mass Spectrometry: data acquisition and analysis

After reduction and alkylation, protein samples were treated with Endoprotease Lys-C (Wako) and Trypsin (Trypsin Gold Mass Spec Grade; Promega). Peptide samples were desalted by OMIX C18 pipette tips (Agilent Technologies) and then analyzed by LC-MS/MS on an LTQ-Orbitrap velos instrument (Thermo Fisher Scientific) connected online to an EASY-nLC system (Thermo Fisher Scientific). Raw mass spectrometry (MS) data from the LTQ-Orbitrap were analyzed using MaxQuant software (Cox & Mann, 2008) version 1.6.10.43, which uses Andromeda search engine (Cox *et al*, 2011). Bioinformatic analysis of the MaxQuant/Andromeda workflow output and the analysis of the abundances of the identified proteins were performed with the Perseus module (Tyanova *et al*, 2016) version 1.6.10.43. Only protein identifications based on a minimum of two peptides were selected for further quantitative studies. After data processing, label-free quantification (LFQ) values from the “proteinGroups.txt” output file of MaxQuant were further analyzed. To distinguish specifically enriched proteins from the background, protein abundances were compared between sample and control groups using the Student’s t-test statistic, and results were visualized as volcano plots (Hubner & Mann, 2011).

### Preparation of puromycin-treated 80S ribosomes from yeast

*S.c*. BY4741 wildtype cells were grown in YP medium with 2% glucose to an OD_600_ of 2.5, then harvested by spinning at 4,400 × *g* for 10 min. Cells were washed first with water, then 1% KCl, then resuspended in 30 ml lysis buffer (20 mM HEPES/KOH pH 7.4, 100 mM KOAc, 7.5 mM Mg(OAc)_2_, 1 mM DTT, 0.5 mM PMSF, complete EDTA-free protease inhibitor mix). Lysis was performed using a Microfluidics M-110L microfluidizer at 15k psi.

The lysate was cleared by centrifugation first at 26,892 × *g* for 15 min., then at 140,531 × *g* for 30 min. 15 ml of cleared lysate were loaded on a layered sucrose cushion consisting of 4 ml 2 M sucrose and 4 ml 1.5 M sucrose (buffer: 20 mM HEPES/KOH pH 7.4, 500 mM KOAc, 5 mM Mg(OAc)_2_, 1 mM DTT, 0.5 mM PMSF) and centrifuged at 246,468 × *g* for 21 h and 15 min.

The pellet containing ribosomal components was resuspended in water and mixed with 2× puromycin buffer (40 mM HEPES pH 7.5, 1 M KOAc, 25 mM Mg(OAc)_2_, 2 mM puromycin, 2 mM DTT, 1 U/ml SUPERase-In RNase Inhibitor (Invitrogen)). The mixture was incubated for 30 min at room temperature, then loaded on 10-40% sucrose density gradients (20 mM HEPES/KOH pH 7.4, 500 mM KOAc, 5 mM Mg(OAc)_2_, 1 mM DTT, 0.5 mM PMSF). Gradients were centrifuged at 20,755 × *g* in an SW 32 Ti rotor (Beckman Coulter) for 20 h. 80S ribosomal fractions were identified using a Biocomp Gradient station *ip* and a Triax Flow cell and were manually collected. Fractions were then pelleted in a TLA110 rotor at 417,200 × g for 45 min and resuspended in storage buffer (20 mM HEPES/KOH pH 7.5, 100 mM KOAc, 5 mM Mg(OAc)_2_, 1 mM DTT). Aliquots were frozen in liquid nitrogen and stored at −80 °C until use.

### Protein expression and purification

#### eIF3j (Hcr1) purification

*Escherichia coli* (*E. coli*) BL21(DE3) cells were transformed with the pTYB2 plasmid containing full length *HCR1* and selected on LB plates containing ampicillin. Cells from a pre-culture were inoculated into 1.5 L of LB medium with ampicillin and cell growth was monitored at 37 °C. At an OD_600_ of 0.6 the cultures were transferred to an ice-water bath and incubated for 20 min. 0.1 mM IPTG was added to induce protein expression and cells were incubated for 15 h at 16 °C while shaking. Cells were harvested by centrifugation at 3,500 × *g* for 10 min and washed with 1% KCl, then resuspended in lysis buffer (20 mM HEPES pH 7.5, 500 mM NaCl). Cells were then pelleted again at 2600 × *g*, frozen in liquid nitrogen and stored at - 80 °C until further use.

Frozen cell pellets were thawed, resuspended in lysis buffer and lysed using a Microfluidics M-110L microfluidizer at 15k psi. The lysate was cleared by centrifugation at 20,000 × *g* for 30 min. Clear lysate fraction was added to 1.5 ml magnetic chitin beads (NEB E8036S) equilibrated in lysis buffer. Binding was performed for 1.5 h at 4 °C on a wheel. Beads were harvested on a magnet and washed once using 5 ml lysis buffer, twice using washing buffer (20 mM HEPES pH 7.4, 1 M NaCl, 1 mM EDTA) and once again using lysis buffer. The protein was then eluted from the beads using 5 ml elution buffer (20 mM HEPES pH 7.4, 500 mM KCl, 50 mM DTT) by incubating on a wheel at 4 °C overnight. A second elution step was performed using the same buffer for one hour after removal of the first elution fraction. Both elution volumes were combined and concentrated using an Amicon Ultra 10k MWCO concentrator. Aliquots of pure eIF3j were flash frozen in liquid nitrogen and stored at −80°C.

#### ABCE1 (Rli1) purification

ABCE1 (Rli1) was overexpressed in *S. cerivisiae* strain WCGα using the pYes2-ABCE1-His_6_ plasmid (kindly provided by R. Green, Department of Molecular Biology and Genetics, Johns Hopkins University School of Medicine) (Shoemaker & Green, 2011). Cells were grown in YP medium lacking uracil and containing 2% galactose, 1% raffinose at 30 °C to mid-log phase and were harvested at a final OD_600_ of 1.0 by centrifugation at 3,500 × *g* for 10 min. Cells were washed once with 1% KCl, pelleted again and resuspended in lysis buffer (75 mM HEPES pH 8.0, 300 mM NaCl, 5mM beta-mercaptoethanol (β-ME), 1% Tween, 20 mM imidazole, 2 mM MgCl_2_, 10% Glycerol). Excess buffer was removed by centrifugation at 2,600 × *g* and the cells were frozen in liquid nitrogen. Frozen cells were ground using a Spex SamplePrep Freezer Mill and the powder stored at −80 °C until further use. The cell powder was thawed and resuspended in lysis Buffer. Cell debris was removed by centrifugation at 47,807.6 × *g* for 30 min and filtered using a 1.6 μm membrane.

ABCE1 was purified first by metal affinity chromatography. Cleared lysate was applied to a HisTrap HP column (GE 5 mL column). The column was washed with 15 colume voulmes (CV) wash buffer (50 mM HEPES pH 8.0, 500 mM NaCl, 5 mM β-ME, 20 mM imidazole, 2 mM MgCl_2_, 10% glycerol) and the protein was eluted with 4 CV over a gradient from 20 mM to 300 mM imidazole. Fractions containing ABCE1 were combined and dialysed against Buffer A (20 mM HEPES pH 7.6, 100 mM KCl, 5 mM β-ME, 0.1 mM EDTA, 10% glycerol, 0.1 mM PMSF) overnight. The sample was diluted to 50 mL and loaded onto a cation exchange column (HiTrap SP 5mL, GE). The column was washed with 6 CV Buffer A and ABCE1 was eluted over gradient from 100 mM to 1 M KCl over 8 CV. ABCE1-containing fractions were concentrated using Amicon® 50k MWCO concentrator before loading onto a gel filtration column (Superdex200) for size exclusion chromatography. The fractions containing ABCE1 were concentrated and aliquots of pure ABCE1 in 20 mM HEPES pH 7.5, 200 mM KCl, 1.5 mM MgCl2, 2 mM β-ME, and 5% glycerol were flash frozen and stored at −80 °C.

#### eIF6 purification

*E. coli* BL21 (DE3) cells were transformed with a p7XC3GH plasmid expressing eIF6 fused to 3C protease cleavage site, GFP and 10-His. Cells were grown on LB medium to mid-log phase (OD_600_ = 0.7-0.8) at 37 °C and induced with 1 mM IPTG at 16 °C for 20 h. Cells were harvested by centrifugation at 4,400 × *g* and 4 °C for 8 min, washed with PBS and resuspended in lysis buffer (20 mM Tris/HCl pH 8.0, 300 mM NaCl, 2 mM β-ME) with 10% glycerol. Resuspended cells were flash-frozen in liquid nitrogen and stored at −80 °C until further use. For purification, frozen cells were thawed and resuspended in lysis buffer without glycerol. Lysis was performed using a Microfluidics M-100L microfluidizer at 15k psi. Crude lysate was cleared by centrifugation at 30,596 × *g* for 20 min. TALON metal affinity resin was equilibrated in lysis buffer and added to the cleared lysate, then incubated at 4 °C for 40 min on a wheel. After collection of the flowthrough, the column was washed using lysis buffer with 10 mM imidazole. Elution was performed by incubating the resin with lysis buffer with 10 mM imidazole and 0.25 *mg/ml* 3C protease for 30 min at 4 °C on a wheel. The elution fraction was concentrated using an Amicon 10k MWCO concentrator and loaded onto a Superdex200 column for size exclusion chromatography using storage buffer (50 mM HEPES/KOH pH 7.5, 500 mM KCl, 2 mM MgCl_2_, 2 mM β-ME). The purified protein in storage buffer was flash-frozen in liquid nitrogen and stored at −80 °C.

Dom34 and Hbs1 were purified as described before (Lee *et al*, 2007).

### Splitting Assays

#### In vitro splitting Assays

Ribosome splitting assays were carried out to test the influence of eIF3j (Hcr1) on the canonical splitting reaction mediated by Dom34, Hbs1, and ABCE1 in yeast. For each reaction, 5 pmol of yeast 80S ribosomes (see above) were mixed with five-fold molar excess of splitting factors Dom34, Hbs1 and ABCE1 as well as the anti-association factor eIF6 under physiological buffer conditions (20 mM HEPES/KOH pH 7.5, 100 mM KOAc, 4 mM Mg(OAc)_2_, 5 mM β-ME, 1 mM ATP, 1 mM GTP). Varying amounts of eIF3j were added to the reactions, ranging from two-fold to twenty-fold molar excess over the 80S ribosomes.

The samples were incubated on ice for 30 min and then loaded on 10-50% sucrose density gradients (20 mM HEPES/KOH pH 7.5, 100 mM KOAc, 5 mM Mg(OAc)_2_, 1 mM DTT, 10-50% (w/v) sucrose). Gradients were spun in an SW 40 Ti rotor (Beckman Coulter) at 202048 × *g* for 4 h and fractionated at a BioComp Gradient Station *ip* using a Triax Flow Cell for UV measurement.

Ribosomal peak fractions were collected manually and from each fraction, proteins were precipitated using 0.015% sodium deoxycholate and 7.2% trichloroacetic acid at 4 °C.

Proteins were separated on a 15% SDS-PAGE gel and visualized using SimplyBlue staining reagent.

#### “Facilitated” splitting assays

“Facilitated” splitting assays were performed to test the association of yeast ABCE1 and eIF3j to ribosomal particles under non-physiological high-salt conditions and in the presence of ATP or the non-hydrolyzable ATP analog AMP-PNP. To induce splitting, purified 80S ribosomes were mixed with ten-fold molar excess of ABCE1 in splitting facilitating buffer (20 mM HEPES/KOH pH 7.4, 500 mM KCl, 1.5 mM MgCl_2_, 1mM DTT). Depending on the experiment, 0.5 mM AMP-PNP or ATP and 10-fold molar excess of eIF3j were added. For the experiments described here, approx. 50 pmol ribosomes in a total reaction volume of 250 μL were used. The samples were incubated for 20 min at 25°C and then cooled down to 4 °C on ice and loaded on 10-50% sucrose density gradients (20 mM HEPES/KOH pH 7.5, 100 mM KOAc, 5 mM Mg(OAc)_2_, 1 mM DTT, 10-50% (w/v) sucrose). All following procedures were carried out as described above for splitting assays.

### Cryo-EM Sample preparation

#### Preparation of native yeast 40S complexes

A BY4741 strain containing genomic TAP-tagged SKI3 and a plasmid overexpressing SKA1 (pCM190) (Zhang *et al*, 2019) was used for generation of the cryo-EM sample.

Yeast cells were grown in synthetic medium lacking uracile (SL -Ura) with 2% glucose at 30 °C to an OD_600_ of 3.0, whereupon the cultures were chilled in ice water. The cells were harvested by centrifugation at 4422 × *g* for 10 min in a Sorvall SLC-6000 rotor, washed with water and resuspended in lysis buffer (20 mM HEPES/KOH pH 7.4, 100 mM KOAc, 5 mM Mg(OAc)_2_, 1 mM DTT, 0.5 mM PMSF, complete EDTA-free protease inhibitor mix). Cells were frozen in liquid nitrogen and ground using a Spex SamplePrep Freezer/Mill.

Frozen cell powder was resuspended in lysis buffer (1:3 w/v) and the lysate was cleared by centrifugation in an SS-34 rotor (Thermo Scientific) at 26891.8 × *g* for 15 min.

Approximately 150 A_260_ absorption units were loaded on a 10-50% sucrose density gradient (buffer composition identical to lysis buffer). Gradients were spun in an SW40 Ti rotor (Beckman Coulter) at 202048 × *g* for 3 h and the 40S peak was harvested manually using a Triax Flow Cell.

Total A_260_ of the collected 40S fraction from yeast lysate was measured and the buffer was exchanged to cryo-EM grid buffer (20 mM HEPES/KOH pH 7.4, 100 mM KOAc, 5 mM Mg(OAc)_2_, 1 mM DTT, 0.5 mM PMSF, complete EDTA-free protease inhibitor mix, 0.05% Nikkol) by three successive rounds of concentration and dilution by a factor of approx. 1:5 using an Amicon Ultra Centrifugal Filter (MWCO 100k) (total dilution factor approx. 1:125). The sample was then concentrated again. The A_260_ was measured as A_260_/ ml = 6.3.

Freshly prepared sample was diluted to approx. 1.25 A_260_ / ml and 3.5 μL were applied to 2 nm pre-coated Quantifoil R3/3 holey carbon support grids and vitrified in liquid ethane using a Vitrobot mark IV (FEI Company, Netherlands). (wait time 45 s, blotting time 2 s).

#### Preparation of native human 40S complexes

Human 40S initiation complexes were found as byproducts in an affinity purification using internally tagged RIOK1 and mutant RIOK1-D324A as bait. In brief, HEK Flp-In 293 T-Rex (Invitrogen) were grown in a 10 cm cell-culture dish to approximately 70% confluency and transfected with 0.5 µg of a pcDNA5/FRT/TO vector containing RIOK1 or RIOK1-D324A and 4.5 µg pOG44 (Invitrogen), using 20 µg polyethyleneimine (PEI). Cells were selected using 150 µg ml-1 hygromycin B (Thermo Scientific) and maintained in DMEM (Thermo Scientific) containing 10% fetal calf serum, 100 µg ml-1 hygromycin B, 10 µg ml-1 blasticidin and 1x penicillin/streptomycin and GlutaMAX (Thermo Scientific). Stable cell lines were subsequently grown in multiple 15 cm cell-culture dishes, protein expression induced with 1.6 µg ml-1 tetracycline and harvested in 0.025% trypsin/EDTA (Thermo Scientific) after 24 h. Cells were washed one in 1x phosphate-buffered saline (PBS) and subsequently pelleted at 1,600 × *g* at 4 °C. Cells were then resuspended in lysis buffer (20 mM HEPES pH 7.6, 150 mM potassium acetate, 5 mM MgCl_2_, 1 mM DTT, 0.5 mM NaF, 0.1 mM Na_3_VO_4_, 1x protease inhibitor (Sigma Aldrich), 0.5% NP-40 substitute) and incubated for 30 min in an over-head rotator at 4 °C, before centrifugation at 4,000 × *g* for 15 min at 4 °C. The cleared lysate was then added to 100 µl of anti-Flag affinity beads (Sigma-Aldrich) and rotated for 2 h at 4 °C. Beads were harvested and 4 times washed with 1 ml wash buffer (20 mM HEPES pH 7.6, 150 mM potassium acetate, 5 mM MgCl_2_, 1 mM DTT, 0.5 mM NaF, 0.1 mM Na3VO4, 1x protease inhibitor (Sigma Aldrich)), before bound complexes were eluted 6 times with 100 µl of 20 mM HEPES pH 7.6, 150 mM potassium acetate, 5 mM MgCl_2_, 1mM DTT, 0.05% Nikkol and 0.2 mg ml-1 3x Flag peptide (Sigma Aldrich). All eluate fractions were combined and concentrated on 300 kDa molecular mass cut-off filters (Sartorius).

3.5 µl of the concentrated sample were applied to glow discharged copper grids with holey carbon support and a 2 nm continuous carbon layer (R3/3, Quantifoil). Grids were blotted in a Vitrobot Mark IV (FEI Company) for 2 s after incubation for 45 s at 4°C and frozen in liquid ethane.

### Cryo-EM data collection and processing

#### Data collection and processing of the yeast 40S complex sample

Cryo EM data were collected on a Titan Krios TEM, using a Falcon II DED at 300 kV, with an electron dose of approx. 2.5 e^-^/Å^2^ per frame for 10 frames. (defocus range of 1.1 to 2.3 µm). The magnified pixel size was 1.084 Å/pixel.

Micrograph stacks collected at the TEM were summed and corrected using MotionCor2 (Zheng *et al*, 2017). Micrograph quality was assessed individually and CTF parameters were estimated using GCTF (Zhang, 2016). Particle picking was performed using Gautomatch (http://www.mrc-lmb.cam.ac.uk/kzhang/) and all further processing was performed using RELION 3.0 (Scheres, 2012; Zivanov *et al*, 2018).

#### Data collection and processing of the human 40s complex sample

Data collection was performed on a Titan Krios at 300 kV, where 7,365 and 4,499 movies were collected for RIOK1-D324A and RIOK1, respectively, at a nominal pixel size of 1.059 Å and at a defocus range from 0.5 to 2.5 µm. Movies were recorded on a K2 Summit direct electron detector using low-dose conditions with 48 frames at approximately 1 e^-^/Å^2^. All frames were gain corrected and subsequently aligned and summed using MotionCor2 (Zheng *et al.*, 2017) and CTF parameters were determined using CTFFIND (Rohou & Grigorieff, 2015) and Gctf (Zhang, 2016). Particles were then picked using Gautomatch (http://www.mrc-lmb.cam.ac.uk/kzhang/). Particle images were extracted in RELION 3.0 (Zivanov *et al.*, 2018) and subjected to reference-free 2D classification. Good particles were selected, 3D refined and classified. Besides the expected pre-40S classes (unpublished), one class containing the initiation complex was obtained in both datasets, comprising approximately 2% (RIOK1-D324A data set) and 8.7% (RIOK1 data set) of the total particle number. The two data sets were subsequently subjected to Bayesian polishing and CTF refinement, combined and further classified extensively as shown in (Suppl. Fig 3). Final reconstructions were then B-factor sharpened with RELION and the local resolution estimated. Where indicated (Suppl Fig 3), local or multi-body refinement was performed.

### Model building and refinement

For rigid body fits and figures Chimera Version 1.13.1 (Pettersen *et al*, 2004) and ChimeraX version 0.91 (Goddard *et al*, 2018) were used. Homology models were created using SWISS-MODEL Repository (Bienert *et al*, 2017; Waterhouse *et al*, 2018).

#### Yeast 43S PIC and 48S IC model

The atomic models PDB 5NDG (Prokhorova *et al*, 2017), 6TB3 (Buschauer *et al*, 2020) (6FYY, and 6FYX (Llacer *et al.*, 2018) containing the models for *S. c.* 40S rRNA, r-proteins and eIFs were fitted as rigid bodies into the cryo-EM maps of the *S.c.* 43S PIC and 48S IC. For the 43S PIC, the 40S rRNA and ribosomal proteins were fitted from PDB 5NDG and eIFs were fitted from PDB 6FYY. For ABCE1, the hybrid semi-open/closed model derived from the human 43S PIC (see below) was fitted into the density. For Hcr1, a homology model was created based on the structure of the human eIF3j dimer (PDB 3BPJ). The C-terminus of protomer 1 was extended by 9 amino acids.

Models for the “mRNA entry position” of the YLC were obtained by fitting the crystal structure of eIF3i/g (PDB 4U1E, Erzberger *et al.*, 2014) to the observed density as a rigid body and matching it to the structure of eIF3b CTD from PDB 6FYY; to obtain the “ES6 position”, the eIF3i-eIF3g moiety bound to the C-terminal helix of eIF3b was rotated by 120 degrees around the Thr697-Asp701 hinge in the CTD of eIF3b as a rigid body.

#### Human 43S PIC

To obtain the atomic model the best resolved maps as obtained after local focused refinement or multi-body refinement (Suppl. Fig. 3 and Suppl. Fig. 5) were used to build the different parts of the *H.s.* 43S PIC. The 40S subunit was fit into maps of 40S body and 40S head obtained from multi-body refinement III (Suppl. Fig. 3) starting with the 40S model (PDB 6G5H, Ameismeier *et al*, 2018). After rigid body fitting, side chains of ribosomal proteins and rRNA were adjusted using Coot (version 0.8.9.2) (Emsley & Cowtan, 2004). Further, the 60S ribosomal protein eL41 was added to the model using PDB 6EK0 (chain h, Natchiar *et al*, 2017). For eIF1A the homology model based on PDB 3J81 (Hussain *et al*, 2014) was fitted and adjusted using the 40S body map. The N-terminal helix bundle of eIF3c (47-149) was built *de novo* into the same map.

The homology model of the crystal structure of the C-terminal part of eIF3d (162-527; PDB 5K4B, Lee *et al.*, 2016) was fitted into the map for the 40S head obtained from multi-body refinement III (Suppl. Fig. 3). The atomic model was only modified in the regions interacting with the 40S head. Similarly, the model for eIF3b (PDB 5K1H, Simonetti *et al.*, 2016) was only adjusted in blades 5 and 6, which contact the 40S body. Here, the best resolved cryo-EM map, obtained by focused classification on the YLC, could be used (Suppl. Fig. 3). Also, the homology model of eIF3i (PDB 5K0Y, Simonetti *et al.*, 2016) and an *α*-helix corresponding to the the C-terminal part of eIF3a were fitted into this map.

The eIF3-PCI-MPN core (including eIF3a, c, e, f, h, k, l, m) was modeled into the two maps of multi-body refinement II (Suppl. Fig. 3) using the human homology model based on PDB 5A5T (des Georges *et al.*, 2015) as starting model. eIF3d-N (2-84) was built *de novo* into the density.

For eIF3j, the unpublished crystal structure of the human eiF3j dimer (PDB 3BPJ) was fitted as rigid body into the density of 43S PIC state II.

Classification of the entire 43S dataset focusing on ABCE1 followed by focused refinement yielded in a well-resolved map, which could be used for model building. The homology model of based on closed-state yeast ABCE1 bound to the 40S (PDB 5LL6, Heuer *et al.*, 2017) was modelled. ATP and ADP were added to the NBSs.

One class obtained by focused classification on the YLC, represents a very stable 43S complex in P_OUT_ state and yielded in a well-resolved map of the TC after focused refinement. The models of tRNA_i_ (PDB 6FEC, Eliseev *et al.*, 2018), eIF2*α* and eIF2*γ* (PDB 6O85, Kenner *et al*, 2019) and the homology models of eIF2*β* and eIF1 (based on PDB 6GSM) were fitted into the map and adjusted using Coot. Further, a stretch of 8 amino acids was modeled into the density adjacent to eIF1 which corresponds to eIF3c.

For the unassigned RRM on top of 18S rRNA h16 and the extra density in the mRNA entry channel as seen in the focused classified maps (Suppl. Fig. 3) we generated a poly-alanine model.

All models were real space refined using Phenix (version 1.17). In order to model the interfaces between the different parts of the structure, maps before and after multi-body refinement were used. Furthermore, neighboring parts were included in the real space refinement using focused cryo-EM maps. The final composite model was subjected to a final refinement using the overall cryo-EM map of state II and state III (Sorting scheme, Table 2). In regions with local resolution lower than 4 Å side chains were not modeled.

### Sequence alignments

In order to quantify the conservation of protein sequences between human and yeast proteins of interest, pairwise alignments were conducted using the T-Coffee implementation at https://toolkit.tuebingen.mpg.de (Notredame *et al*, 2000; Zimmermann *et al*, 2018) and visualized using JalView (Waterhouse *et al*, 2009). Multiple sequence alignments of the conserved elements of the eIF3c N-terminus were created using MAFFT (Katoh *et al*, 2019).

## Supplementary Figures

**Supplementary Figure 1:**
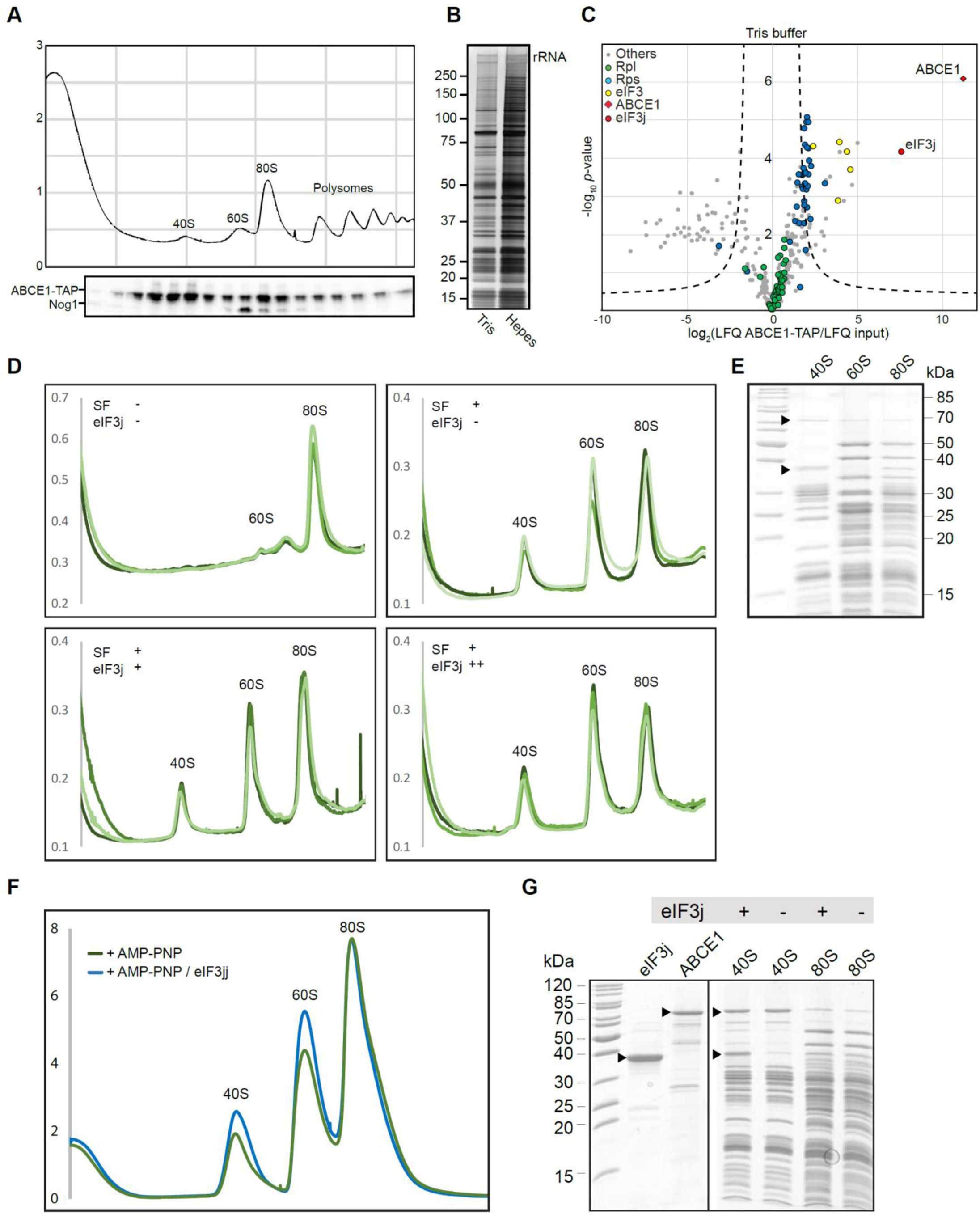
Enrichment of ABCE1 and eIF3j on 40S complexes and assessment of their role in splitting of 80S ribosomes. (A) Total cellular extracts from yeast cells expressing ABCE1-TAP were separated on a sucrose gradient (10–50%) by ultracentrifugation. Proteins of each fraction were analyzed by Western blot using a PAP antibody for the detection of the ABCE1-TAP fusion protein and anti-Nog1 antibody to mark the 60S fraction. (B) Silver stained NuPAGE gel showing elution from affinity purification using ABCE1-TAP performed in Tris or Hepes buffer (see methods for details). (C) Volcano plot showing the fold enrichment of proteins in the elution fraction from the ABCE1-TAP purification in Tris buffer followed by mass spectrometry analysis (LC-MS/MS). The enrichment was calculated relative to an “input” corresponding to an aliquot of the ABCE1-TAP cell lysate used for the affinity purification. It is represented, on the x-axis, as log_2_(LFQ ABCE1-TAP/LFQ input) where LFQ stands for label-free quantification. The y-axis represents the P-value distribution (-log_10_*-p*-value) calculated using the Student’s t-test for all identified proteins represented by a circle. Proteins above the curved lines show a statistically significant enrichment according to the t-test value. The assay was performed in triplicates. (D) UV-profiles from *in vitro* splitting reaction triplicates with and without splitting factors (SF; ABCE1, Dom34, Hbs1 and eIF6) and eIF3j. Samples were separated on a sucrose gradient (10–50%) by ultracentrifugation. (E) SDS-PAGE of the 40S, 60S and 80S peaks obtained from the i*n vitro* splitting experiment (D) containing SFs and high amounts of eIF3j. (F) UV-profiles from *in vitro* “facilitated” splitting reactions. Samples were separated on a sucrose gradient (10–50%) by ultracentrifugation. (G) SDS-PAGE of the input factors (eIF3j and ABCE1) as well as 40S and 80S peaks from the “facilitated” splitting experiment.

**Supplementary Figure 2:**
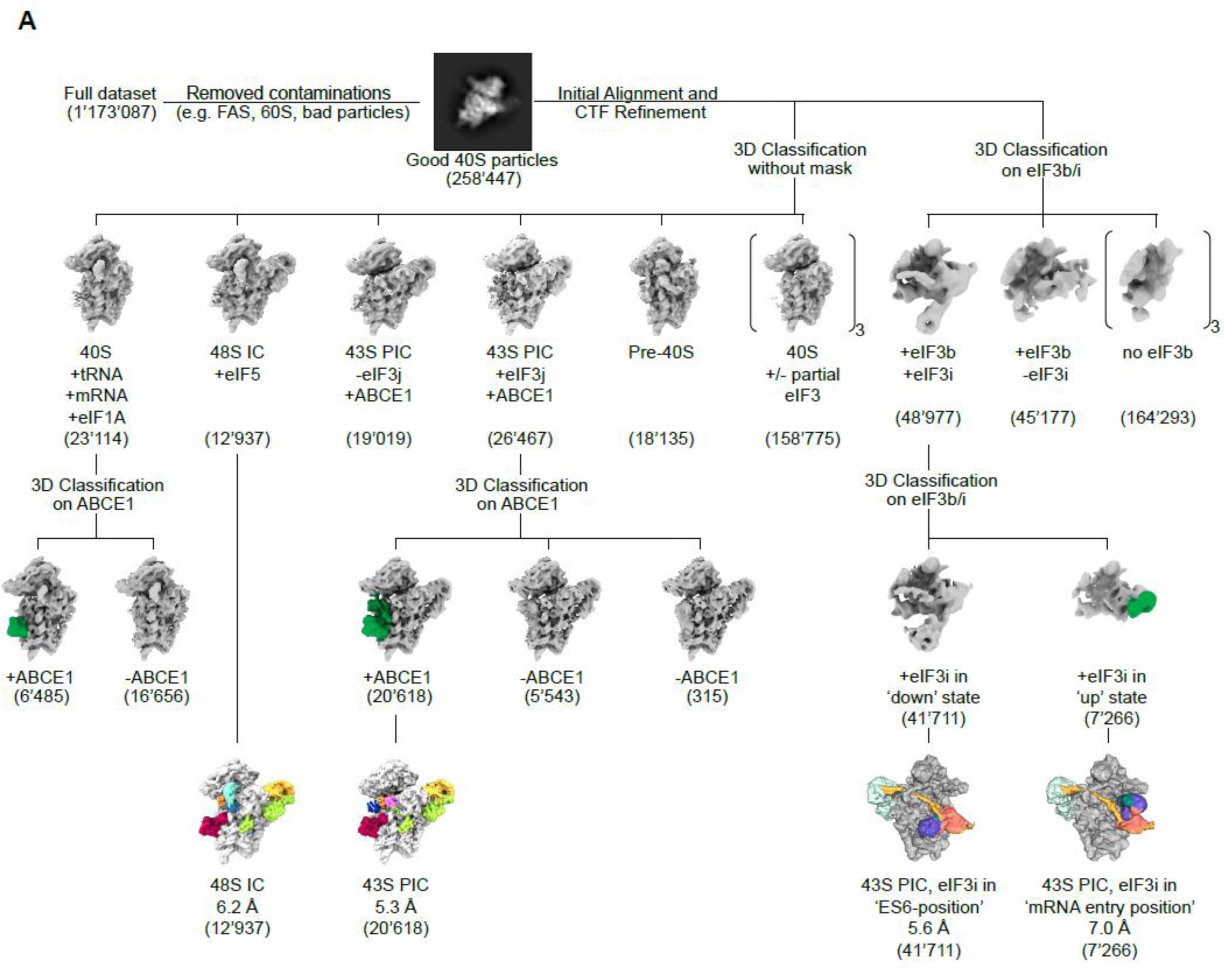
Sorting scheme for the yeast native small 40S sample. After 2D classification of the full dataset, approximately 260.000 particles representing 40S subunits were selected. By 3D classification into six classes, 29% of the particles were unambiguously identified as (pre-) initiation complexes, amongst them a 43S PIC class containing eIF1, eIF1A, eIF3 and ABCE1 with stably bound eIF3j and one class representing a partial 48S initiation complex containing eIF1A, eIF3, eIF5, the TC, and ABCE1. The 48S IC was refined to 6.2 Å and the eIF3j-containing 43S was sub-classified for ABCE1, yielding a 5.3 Å reconstruction of the 43S PIC. In total, 62% of the found initiation-factor-containing 40S particles contained ABCE1 after focused classification for ABCE1 presence. Independently, focused classification on the particles containing the eIF3b/i/g module was performed to assess conformational distribution of eIF3i/g with respect to eIF3b. 85% of the particles contained eIF3i in the “ES6-position”, while in 15% of particles, eIF3i was in the “mRNA entry-position”.

**Supplementary Figure 3:**
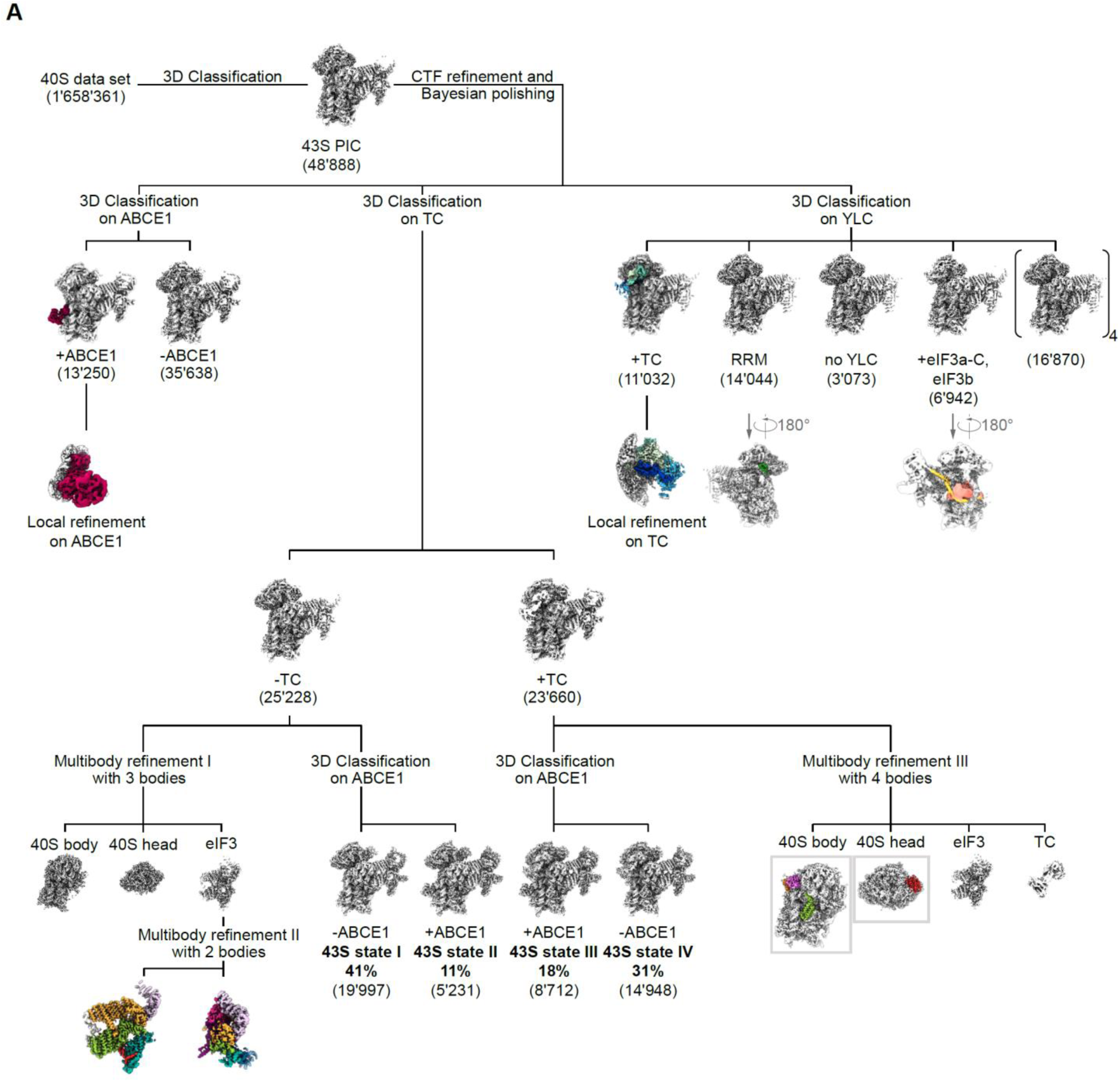
Sorting scheme for the human native small 40S sample. The data set was first classified for presence of initiation factors (see Materials and Methods). 2.9% of all particles contained the eIF3 PCI-MPN core at the back side of the 40S and partial densities for the YLC at the mRNA entry site, the TC in the ISS, and ABCE1. Focused classifications were performed using a binary mask with soft edges to obtain homogenous populations of 43S complexes. Focused classification on the TC yielded two classes with and without the TC that also differed in the conformation of the 40S head (closed latch without TC and P_OUT_ state). These two classes were sub-classified focusing on ABCE1, yielding the four main classes shown in Fig. 2. To obtain the highest possible resolution, we independently performed multi-body refinements on TC-bound and - unbound classes. The TC-lacking class containing the eIF3-PCI-MPN core was refined in two steps: first, the 43S was divided into three bodies (40S body, 40S head and eIF3; multi-body refinement I). Then the body containing the eIF3-PCI-MPN core was re-centered, the 40S SSU signal subtracted and a multi-body refinement with two bodies was performed (multi-body refinement II) that were used for model building. Particles containing the TC were subjected to multi-body refinement III with four bodies (40S body and head, eIF3-PCI-MPN core and TC) yielding well-resolved densities for the eIF3c-NTD on the body and eIF3d on the head used for model building. Classification of the entire 43S data set focusing on ABCE1 followed by focused refinement yielded a well-resolved map from 27% ABCE1-containing particles used for model building. Focused classification on the YLC revealed various compositional and conformational states. One class represented a very stable complete 43S complex in P_OUT_ state and focused refinement yielded a well-resolved map of the TC used for model building. A focused refinement resulted in the best resolved map for the TC, which could be used for model building; two other classes were enriched in stably bound YLC, one showing the clear connection between the PCI-MPN core and the YLC and one with a well-resolved density for a RRM adjacent to the mRNA entry site and density in the mRNA channel.

**Supplementary Figure 4:**
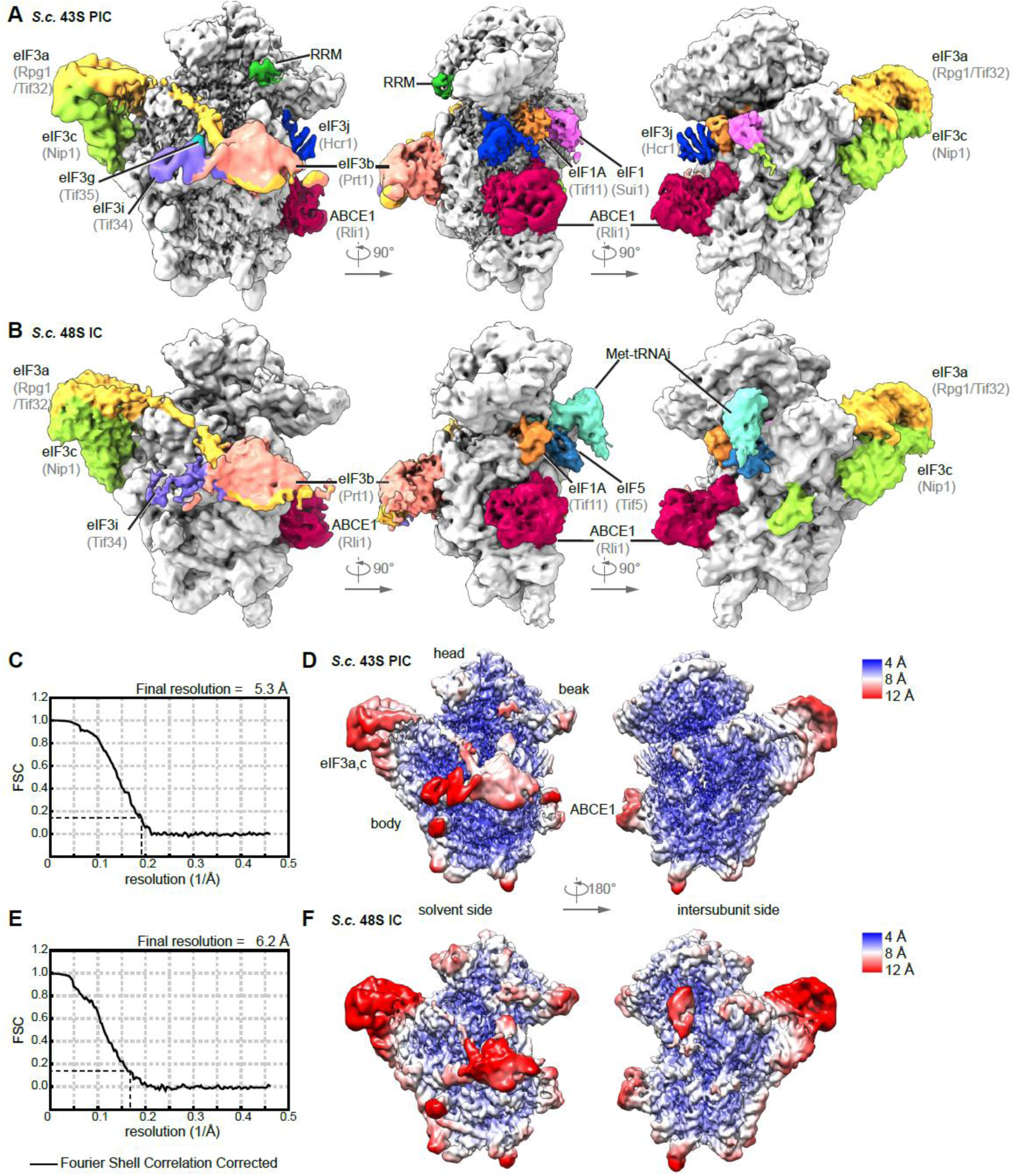
Overview and resolution of the yeast 43S PIC and 48S IC. (A) and (B) Three views on the 3D reconstructions of the yeast 43S PIC (A) and the 48S IC (B) low-pass filtered according to local resolution. (C) Gold standard Fourier Shell Correlation (FSC) curve and (D) 3D reconstruction the yeast 43S PIC colored and filtered according to local resolution. (E) Gold standard Fourier Shell Correlation (FSC) curve and (F) 3D reconstruction the yeast 48S IC colored and low-pass filtered according to local resolution.

**Supplementary Figure 5:**
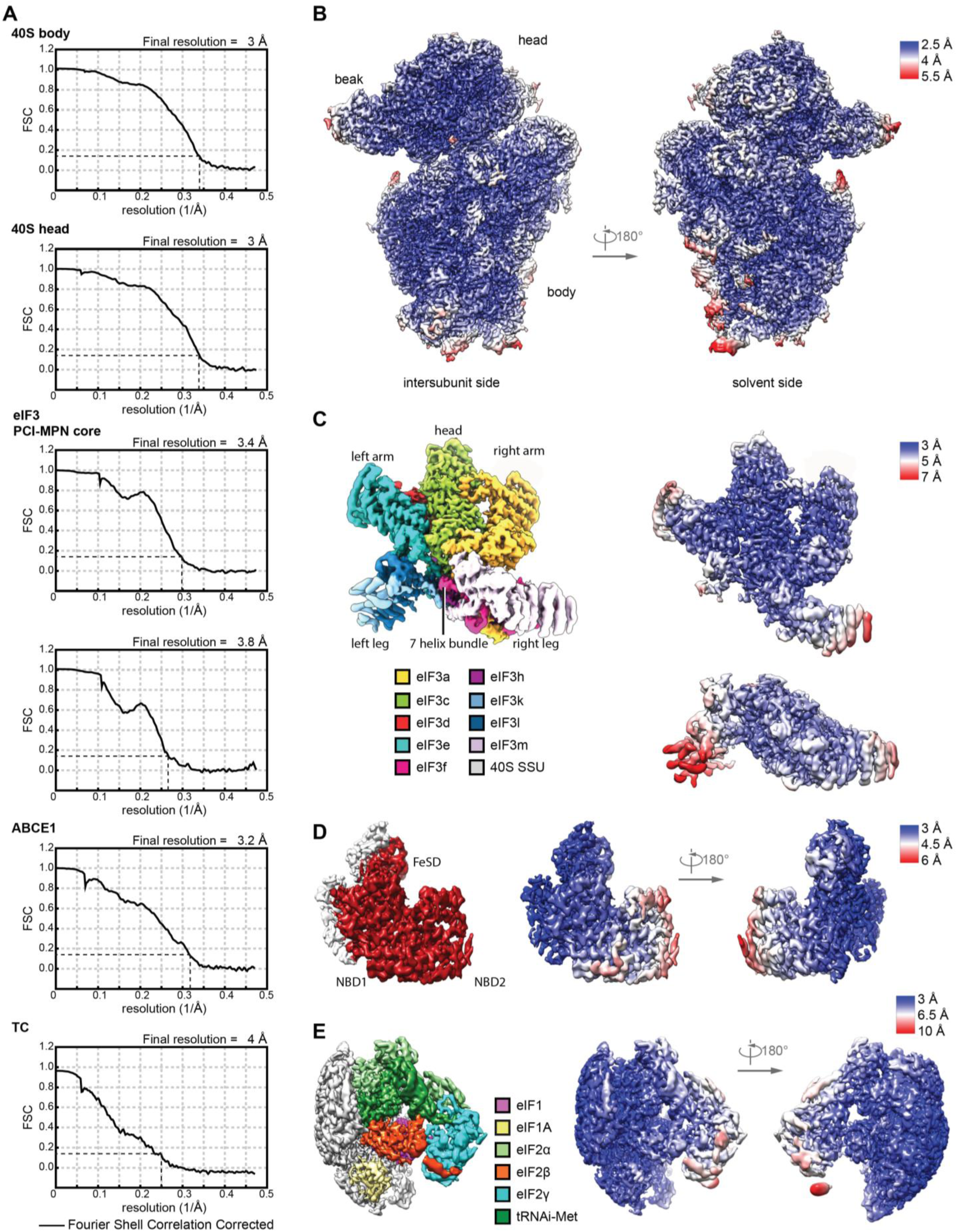
Resolution and model fitting of the human 43S PIC. (A) Gold standard Fourier Shell Correlation (FSC) curve for individual bodies in multi-body refinements (40S head including eIF3d, 40S body including eIF3c-NTD and eIF1A, two bodies of the eIF3 PCI-MPN core) and focused refinements using soft binary masks (ABCE1 and the TC). (B) Composite map of 40S head and body after multi-body refinement colored and low-pass filtered according to local resolution. (C) Composite map of the eIF3 MPN-PCI core (left) after multi-body refinement with two bodies (right), filtered according to local resolution. Structural hallmarks are color coded in the composite map and the two bodies are colored according to local resolution. (D) Focused refined map of the masked region containing ABCE1, filtered according to local resolution and colored for ABCE1 (left). Two views are shown colored according to local resolution (right). (E) Same as in D for eIF1, eIF1A, and the TC.

**Supplementary Figure 6:**
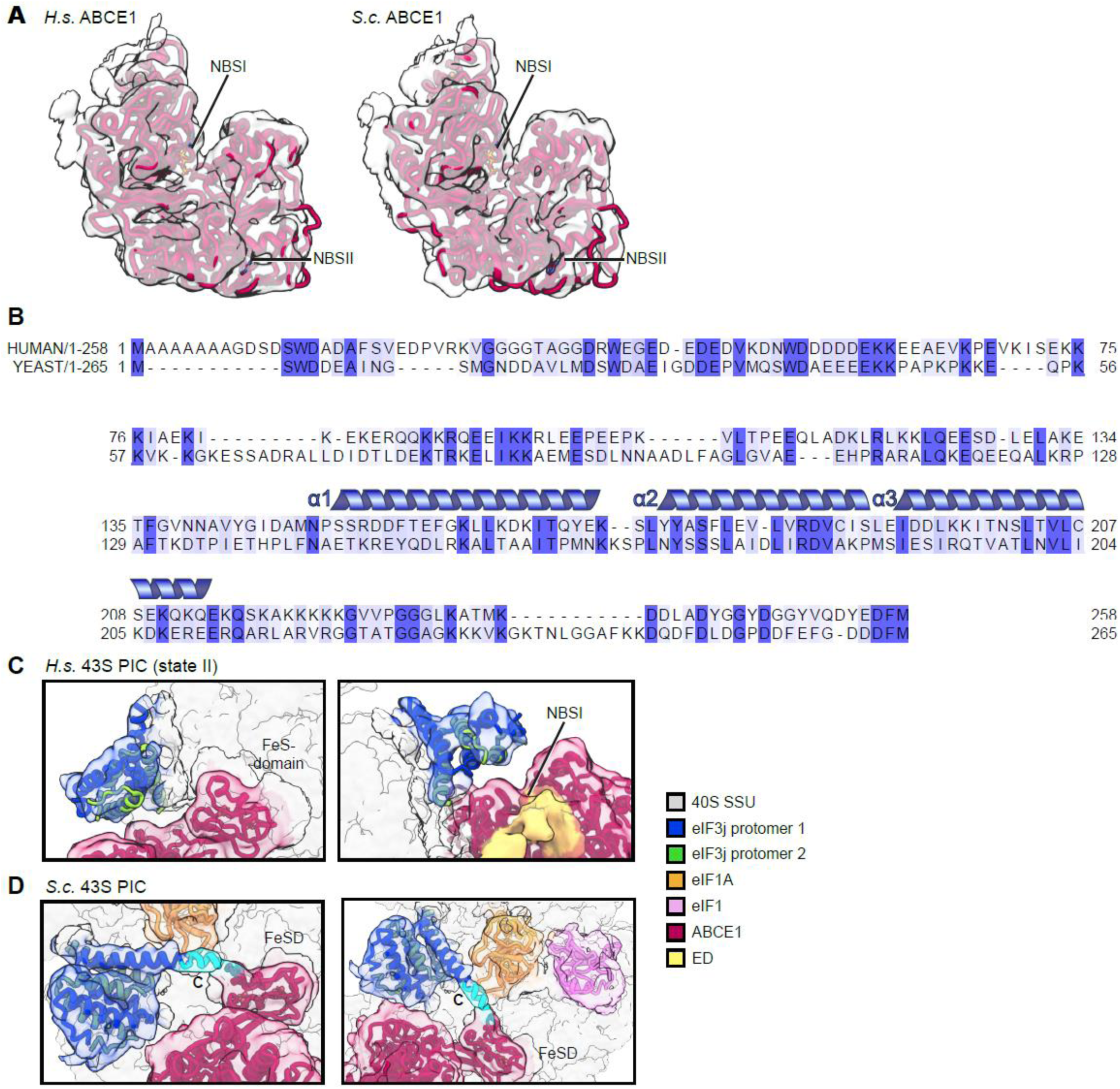
Density fits of ABCE1 and alignment, model and ribosome binding of human and yeast eIF3j. (A) Model for ABCE1 fit into low-pass filtered density to demonstrate the hybrid semi-open/closed conformation of ABCE1 in native yeast and human 43S PICs. (B) Alignment between *H. s.* and *S. c.* eIF3j shows 24,6% identity and 53,1% similarity for the full-length protein. For the sequence (three *α*-helices) present in the human X-ray structure (PDB 3BPJ from residues 144-213 in protomer 1 and 144-216 in protomer 2) identity/similarity is 32.4%/66.2%, corroborating the reliability of the yeast homology model. (C, D) Fits of the human eIF3j crystal structure and the yeast homology model into the corresponding density. In the yeast model, the C-terminal helix of protomer 1 was extended by 9 residues (shown in cyan).

**Supplementary Figure 7:**
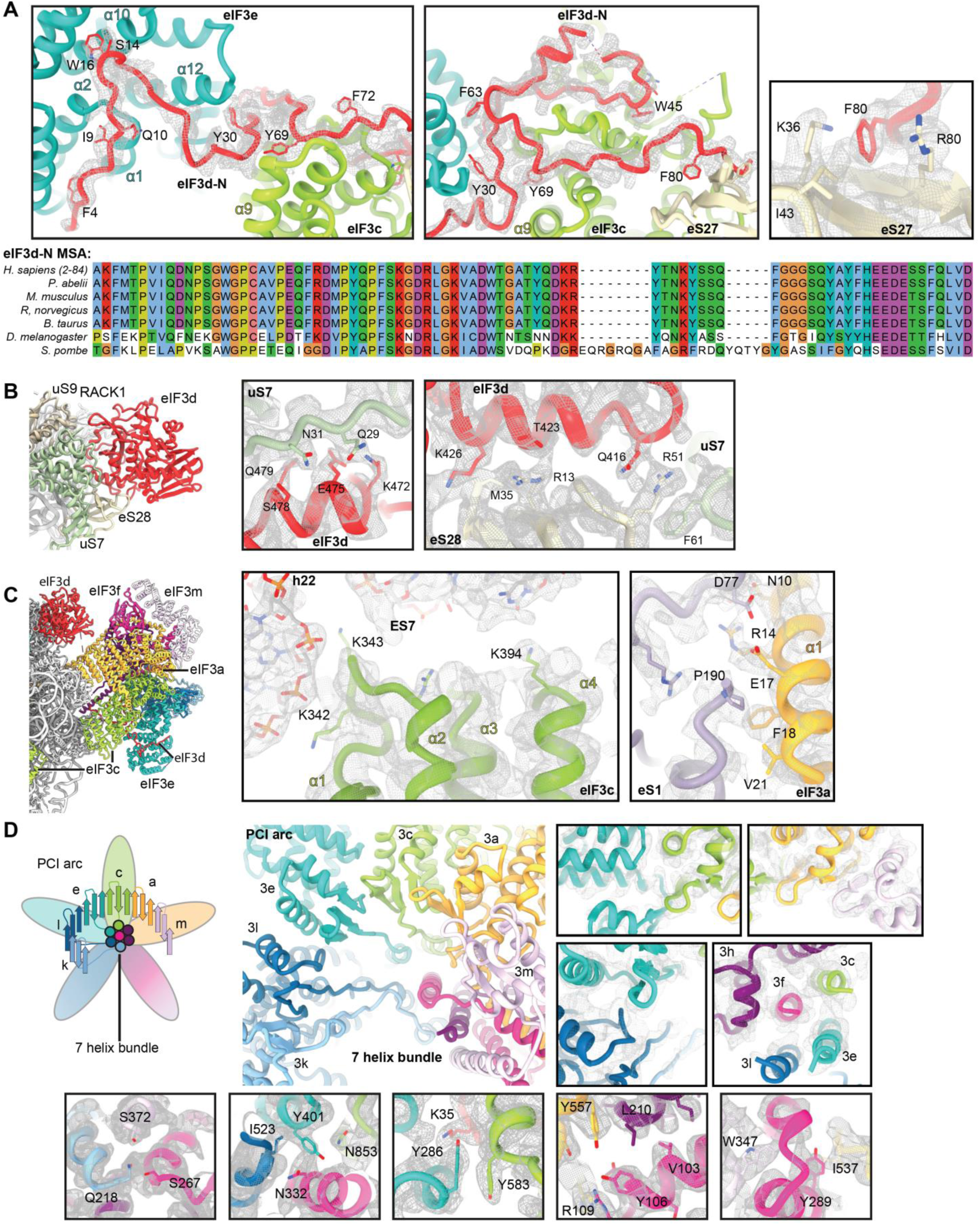
Molecular interactions of eIF3d and the PCI-MPN core in the 43S PIC shown with density. (A) Interactions of the eIF3d N-terminal tail with the PCI-MPN core: interactions of the ultimate eIF3d N-terminus (Phe4-Pro18) to eIF3e are established *via* Phe4 (to Tyr32 in loop between PCI helices *α*2 and *α*3), Gln10 (to His12 in *α*1), Ile9 and Asp11 (to Arg16 in *α*2), Ser14 (to Asn164 *α*9 and Phe132 in *α*7), Trp16 (to Gly171 in *α*10), Gly17 (to Trp170 in *α*10). eIF3d residues 25-36 are bridging eIF3e and eIF3c. Residues involved are Tyr30 (to Leu208 in eIF3e PCI helix *α*12), Phe33 (to the peptide backbone of Leu590 in eIF3c *α*11) and Lys35 (to Gln283 in eIF3e *α*16). eIF3c specific interactions are established by Leu39 (to Gln595), Trp45 (Pro603, Ile607 and Glu666) and Thr46 (to Arg641) (see Table 3). For clarity, only density for eIF3d is shown (grey transparent mesh). (B) Interactions of the eIF3d C-terminal domain with the 40S head and zoomed views of uS7 and eS28 interaction sites shown with density. (C) eIF3a and eIF3c interactions to the 40S. (D) PCI arc of the PCI-MPN core; zoomed views highlighting fits of the eIF3 PCI arc into the refined density and interactions between the subunits in the PCI-MPN core.

**Supplementary Figure 8:**
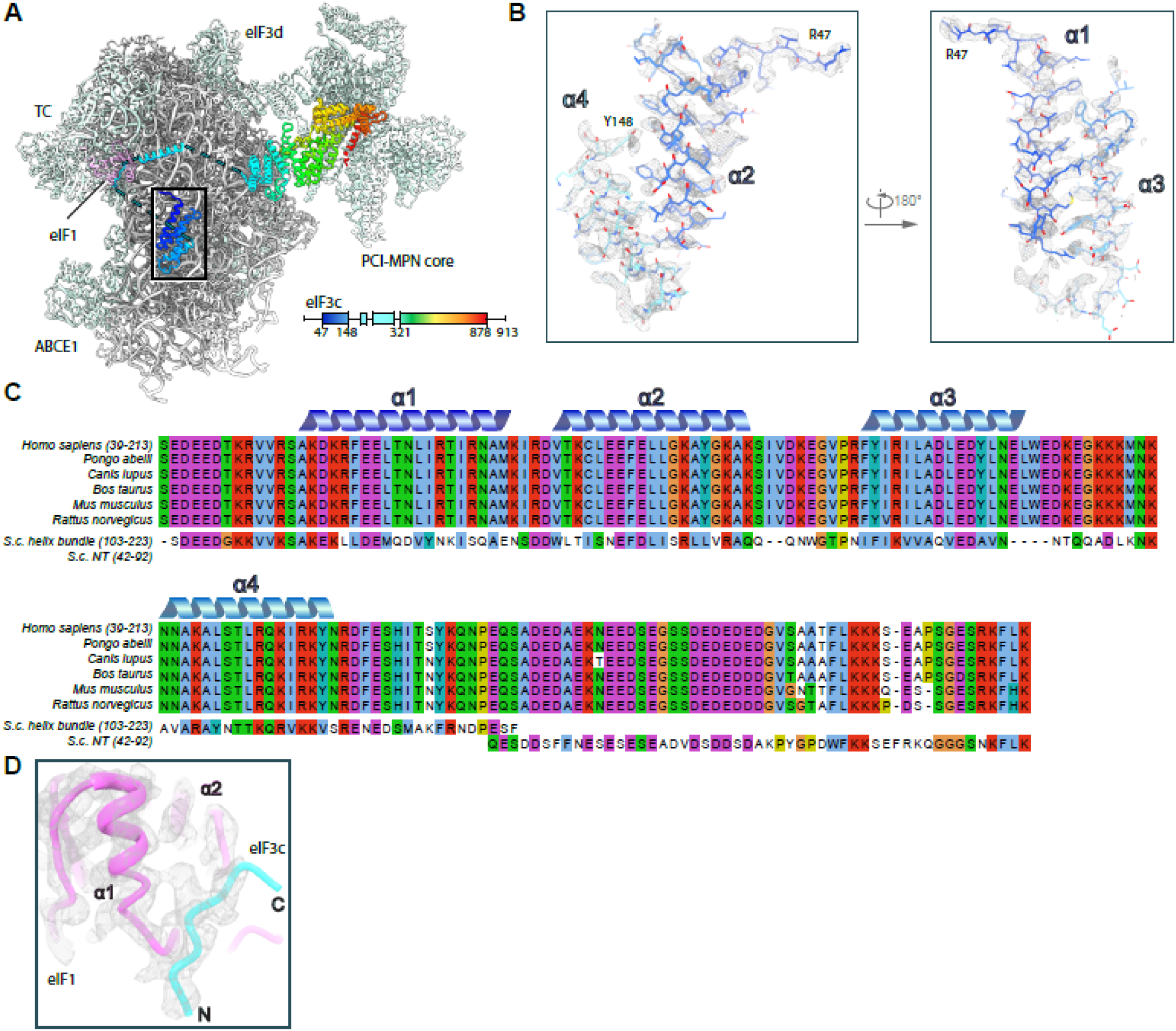
Model fitting and sequence alignment of the eIF3c-NTD. (A) Overview of the TC-containing human 43S PIC as shown in Figure 5 and scheme indicating the parts of eIF3c modeled. (B) Zoomed views highlighting fits of the eIF3c-NTD 4-helix bundle into the refined density (grey transparent mesh). The N- and C-terminal residues are marked. (C) MSA of the conserved N-terminal region of eIF3c in mammals (Ser39-Lys213 in *H.s.*), aligned with segments of the NT from *S.c..* The 4-helix bundle shows 31.1/67.2% sequence identity/similarity, and the eIF1-interacting stretch present in the N-terminus of *S.c.* eIF3c (Gln42-Lys92) shows 32.0/56.0% sequence identity/similarity with a mammalia-specific insert C-terminal of the conserved 4-helix bundle. (D) Zoomed view showing the fit of the eIF1-interacting stretch of eIF3c into the cryo-EM density.

**Supplementary Figure 9:**
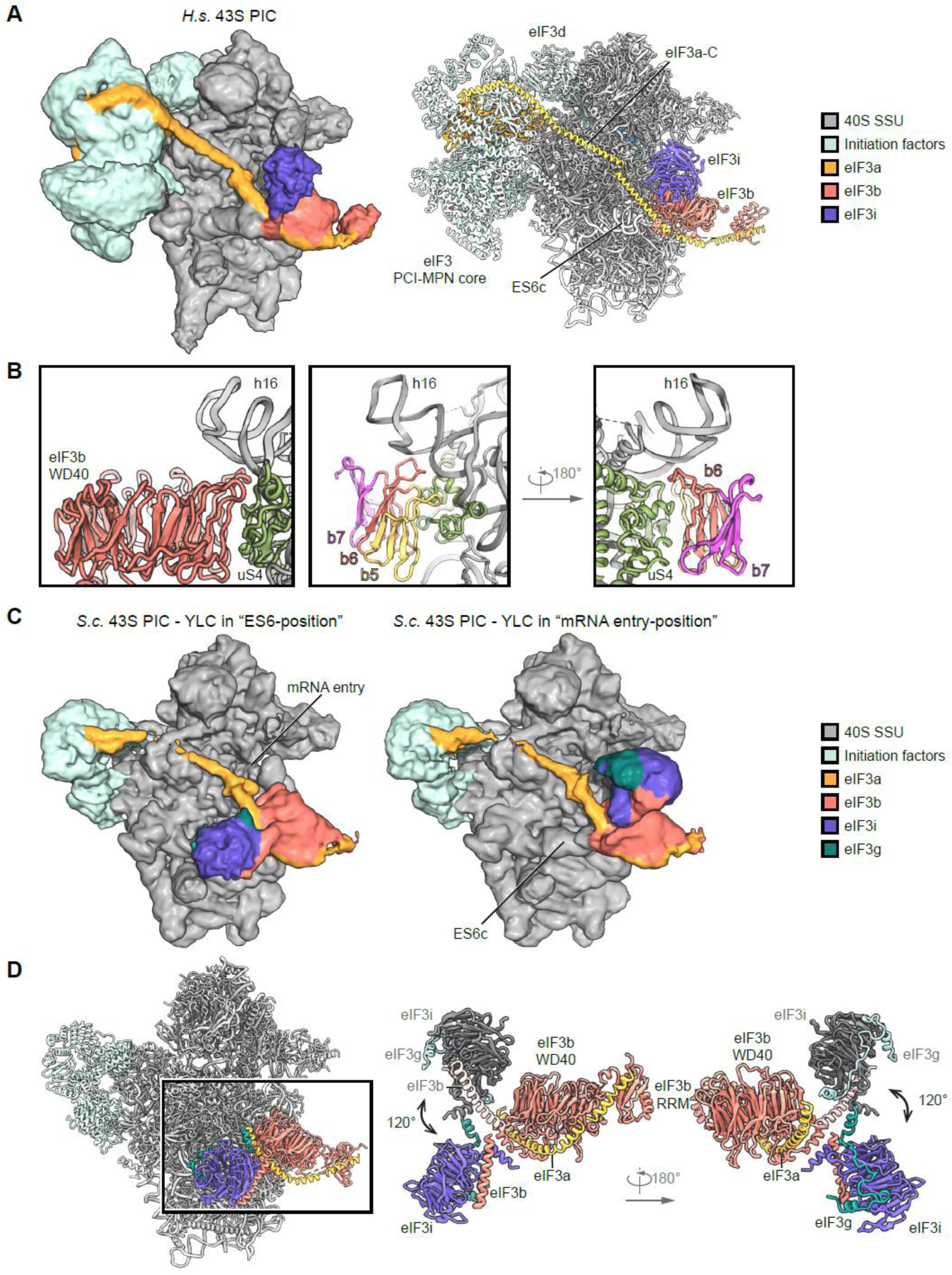
Position of the eIF3 YLC and the eIF3a-CTD in yeast and human 43S PICs. (A) Cryo-EM map of the human 43S PIC class obtained after local classification on the YLC (see Suppl. Fig. 3), low-pass filtered at 6 Å (left). Composite model of the human 43S PIC as in Fig. 2 (right). The view focuses on the mRNA entry side of the 40S, showing the YLC and the rod-like eIF3a density representing its C-terminus spanning from the back side of the 40S to the YLC. (B) Interactions of eIF3b with the ribosome; models for eIF3g and eIF3a not shown. Two different views show only WD40 blades (b) 5 to 7 of eIF3b. In *β* −strand D5 (nomenclature refers to Liu *et al*, 2014) Arg505, Arg507 and Leu509 and in *β* −strand D6 Val558, Glu560 are facing towards uS4. The loop between B5-C5 (485-490) interacts with the rRNA backbone of the h16-h17 junction and uS4 (Tyr 165) and the loop between D5 and A6 (especially Phe510) interacts with Lys121 of uS4. h16 is contacted *via* the loop B6-C6 (res 532-541) *via* backbone interactions. (C) Two different states of the yeast YLC obtained after focused classification. In one state (“ES6-position”) the eIF3g-eIF3i module bound to the eIF3b most C-terminal helix is facing towards expansion segment ES6c, in the other state (“mRNA entry-position”) it faces towards the mRNA entry, similar as in the human 43S PIC and as described previously (Erzberger *et al.*, 2014; Llacer *et al.*, 2018). 85% of the particles contained eIF3i in the ES6-position and 15% of particles in the mRNA entry-position (see Suppl. Fig. 2). (D) Molecular model of the yeast 43S PIC with the YLC in (left) and overlay of the two positions. In the overlay the mRNA entry-position eIF3i is colored gray and eIF3g light blue and the eIF3b C-terminal helix white. The loop between the two most C-terminal helices of the eIF3b CTD (Thr697-Asp701) serves as a hinge for rotation.

**Supplementary Figure 10:**
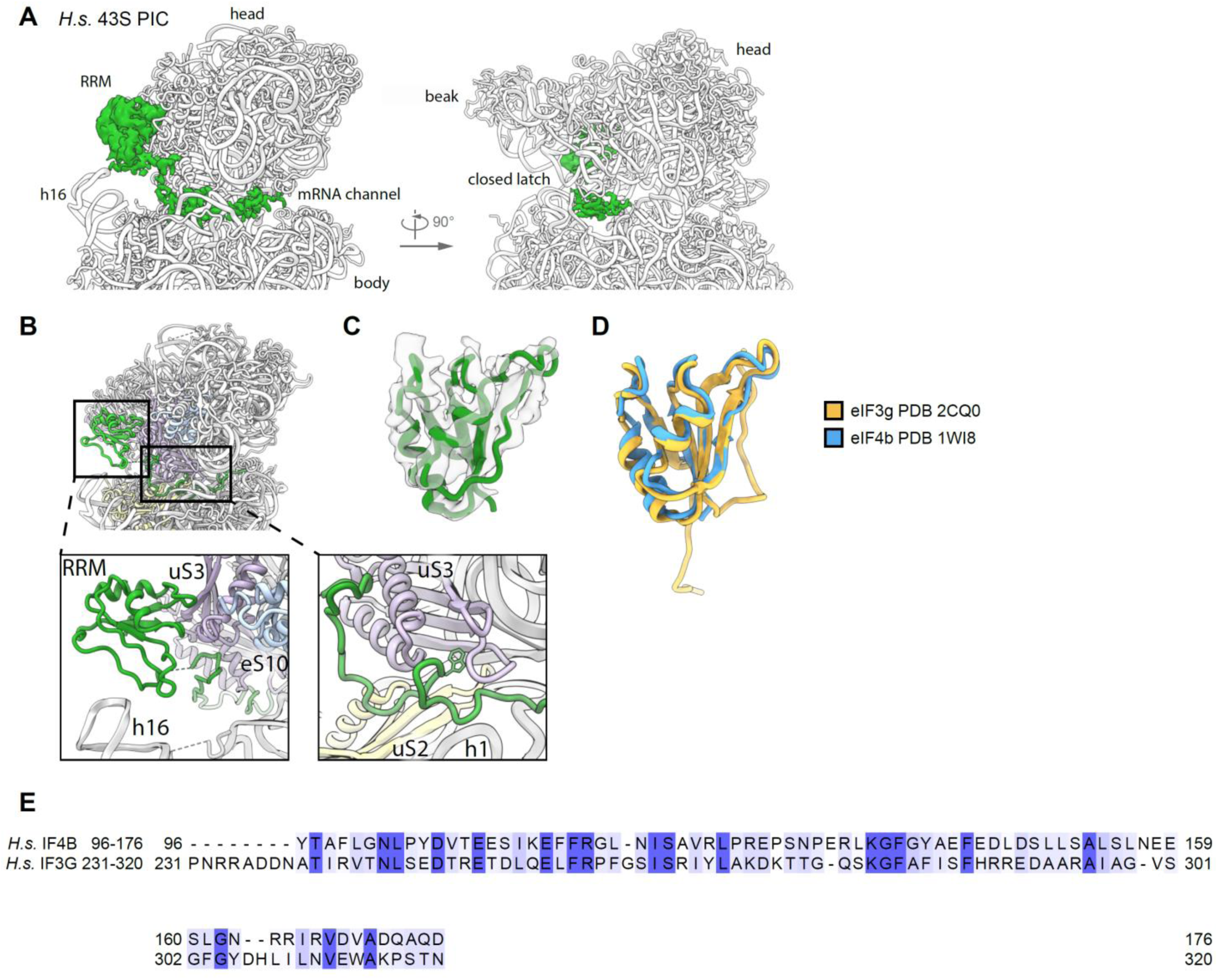
Position of the eIF3g or eIF4b RRM and density in the mRNA channel. (A) Zoomed views on the mRNA channel as viewed from the ISS focusing on an extra density (green) on top of rRNA h16 and inside the mRNA channel. The isolated density is low-pass filtered according to local resolution. (B) Overview and zoomed views on the poly-alanine model for an RRM on top of h16 and for the density in the mRNA channel in context of the 43S PIC. Interacting r-proteins and rRNA and the clearly visible tryptophan residue interacting with uS3 in the mRNA entry channel are highlighted. (C) Model for a typical RRM-fold fitted into the corresponding isolated density. (D) Overlay of the RRMs of eIF4b and eIF3g. (E) Sequence alignment of the RRM of human eIF4b and eIF3g, which shows 21.7/55.4% sequence identity/similarity.

**Supplementary Figure 11:**
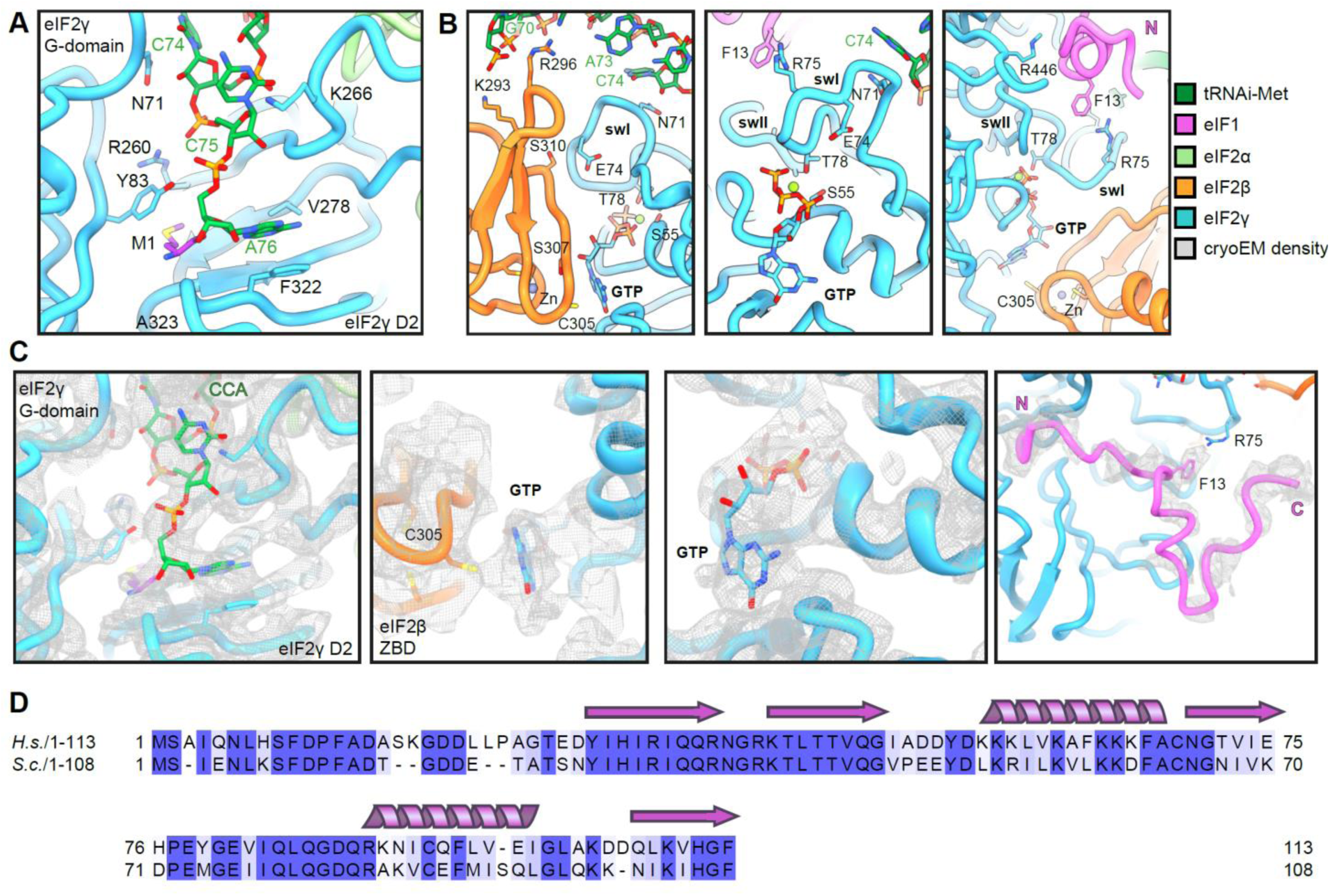
Molecular interactions of the TC in the complete human 43S PIC. View focusing on (A) the interactions of methionylated tRNA_i_ with eIF2γ, (B) the ZBD of eIF2β packing upon the nucleotide binding pocket of eIF2γ and binding to tRNA_i_ (left) and the switch loops (sw) of eIF2γ contacting the *de novo* built eIF1 N-terminal tail (middle and right). (C) Zoomed views of fits of the TC model into the cryo-EM map. Highlighted are the CCA-end of tRNA_i_ bound to eIF2γ, the guanine base lock-up by the eIF2β ZBD, the GTP in the eIF2γ nucleotide binding pocket and the *de novo* built N-terninal tail (res 4-30) of eIF1. (D) Sequence alignment between the yeast and human eIF1 shows a sequence identity of 61.1% and a sequence similarity of 87.0% (N-terminus of eIF1 (4-30) shows 55.6/74.1% sequence identity/similarity) indicating a high degree of conservation.

